# Unfolded Von Willebrand Factor Binds Protein S and Reduces Anticoagulant Activity

**DOI:** 10.1101/2024.02.08.579463

**Authors:** Martha M.S. Sim, Molly Y. Mollica, Hammodah R. Alfar, Melissa Hollifield, Dominic W. Chung, Xiaoyun Fu, Siva Gandhapudi, Daniëlle M. Coenen, Kanakanagavalli Shravani Prakhya, Dlovan F. D Mahmood, Meenakshi Banerjee, Chi Peng, Xian Li, Alice C. Thornton, James Z. Porterfield, Jamie L. Sturgill, Gail A. Sievert, Marietta Barton-Baxter, Ze Zheng, Kenneth S. Campbell, Jerold G. Woodward, José A. López, Sidney W. Whiteheart, Beth A. Garvy, Jeremy P. Wood

## Abstract

Protein S (PS), the critical plasma cofactor for the anticoagulants tissue factor (TF) pathway inhibitor (TFPI) and activated protein C (APC), circulates in two functionally distinct pools: free (anticoagulant) or bound to complement component 4b-binding protein (C4BP) (anti-inflammatory). Acquired free PS deficiency is detected in several viral infections, but its cause is unclear. Here, we identified a shear-dependent interaction between PS and von Willebrand Factor (VWF) by mass spectrometry. Consistently, plasma PS and VWF comigrated in both native and agarose gel electrophoresis. The PS/VWF interaction was blocked by TFPI but not APC, suggesting an interaction with the C-terminal sex hormone binding globulin (SHBG) region of PS. Microfluidic systems, mimicking arterial laminar flow or disrupted turbulent flow, demonstrated that PS stably binds VWF as VWF unfolds under turbulent flow. PS/VWF complexes also localized to platelet thrombi under laminar arterial flow. In thrombin generation-based assays, shearing plasma decreased PS activity, an effect not seen in the absence of VWF. Finally, free PS deficiency in COVID-19 patients, measured using an antibody that binds near the C4BP binding site in SHBG, correlated with changes in VWF, but not C4BP, and with thrombin generation. Our data suggest that PS binds to a shear-exposed site on VWF, thus sequestering free PS and decreasing its anticoagulant activity, which would account for the increased thrombin generation potential. As many viral infections present with free PS deficiency, elevated circulating VWF, and increased vascular shear, we propose that the PS/VWF interaction reported here is a likely contributor to virus-associated thrombotic risk.

**Key Points:**

- Von Willebrand Factor (VWF) binds Protein S (PS) in a shear-dependent manner, reducing the free PS pool and its anticoagulant activity.
- The PS/VWF complex forms under turbulent flow conditions, is stable in whole blood, and localizes to growing platelet thrombi.

## Introduction

Hemostasis is a tightly regulated balance of procoagulant proteins, which promote the activation of thrombin and formation of a fibrin clot, and anti-coagulant proteins, which terminate this process.^1^ Thrombosis, occurs due to a dysregulation of this balance, caused by increased procoagulant activity, decreased anticoagulant activity, or both. Protein S (PS) is a critical anticoagulant protein, present in plasma and platelet α-granules,^2^ deficiency of which is associated with increased thrombotic risk.^3^ Plasma PS exists in two pools. Approximately 40% is considered “free PS” and has anticoagulant activity.^4^ It is a cofactor for activated protein C (APC), which degrades factors Va and VIIIa,^5^ and tissue factor pathway inhibitor-alpha (TFPIα), which inhibits factors VIIa and Xa.^6^ PS also directly inhibits factor IXa.^7^ The remaining ∼60% of plasma PS is bound to the β-chain of complement component C4b-binding protein (C4BP-β).^4^ C4BP stabilizes PS, preventing its clearance,^8^ and blocks most anticoagulant activity.^9^ Acquired PS deficiency is a common complication of severe viral infections, including human immunodeficiency virus,^10,11^ varicella,^12^ dengue,^13^ and severe acute respiratory syndrome-coronavirus-2 (SARS-CoV-2),^14–19^ all of which are associated with an increased risk of thrombosis.

There are two broad categories of acquired PS deficiency: loss of total or free PS. While the causes of total PS deficiency are varied (*e.g.*, decreased synthesis, increased consumption, or degradation^18^), the causes for loss of free PS are unclear. Because total PS is unchanged, free PS deficiency is thought to indicate the presence or increase of a PS-binding protein. The most widely studied example of this is C4BP, which co-precipitates with PS with polyethylene glycol (PEG).^20^ However, while C4BP does increase during inflammation, the change is primarily C4BP α-chain, as opposed to the PS-binding β-chain.^21^

Here, we focus on identifying other PS binding partners in plasma that could cause PS deficiencies. We demonstrate that PS interacts with von Willebrand Factor (VWF) in a shear-dependent manner. Sheared VWF blocks the detection of free PS and reduces PS anticoagulant activity in human plasma. Additionally, the PS/VWF complex forms as VWF unfolds under flow, is stable under arterial flow conditions, and localizes to growing platelet thrombi. We finally show the potential clinical relevance of our observations in a cohort of SARS-CoV-2+ patients, which presented with free PS deficiency that correlated with increased VWF and thrombin generation. Based on our data, we propose a mechanism of free PS deficiency under inflammatory conditions, where the increased presence of sheared VWF sequesters PS and thereby limits its anticoagulant activity.

## Methods

### Data availability

All data included in this study are available upon request from the corresponding author.

### Study population and sample isolation

Human subject studies were approved by the Institutional Review Board of the University of Kentucky (UK) and the procedures followed were per the Helsinki Declaration. Written informed consent was received from subjects before participation. Blood samples were collected from adults: SARS-CoV-2 negative controls (n=38, 56% male, 44% female, age 59.9±14.1 y) and hospitalized SARS-CoV-2+ (COVID-19) patients (n=30, 69% male, 31% female, age 61.5±14.2 y). Subjects were enrolled between April 2020 and January 2021 through the Kentucky Clinic and the University of Kentucky Center for Clinical and Translational Science. Blood was collected and plasma was isolated as described.^25^ For some experiments, samples were also obtained from SARS-CoV-2 positive outpatients (patients with mild or no symptoms, post-quarantine; n=5, 40% male, 60% female, age 48.4±15.5 y).

### Statistics

Statistical analyses were performed using GraphPad Prism version 9.5.1. Grouped data were analyzed by non-parametric Mann-Whitney tests (two groups) or Kruskal-Wallis followed by Dunn’s multiple comparisons tests (more than two groups), and correlation coefficients were calculated by the method of Spearman, unless otherwise noted in figure legends.

**Additional methods and materials can be found in supplemental data**

## Results

### VWF interacts with PS in a shear-dependent manner

Several disease states present with free PS deficiency, and many have a concomitant increase in VWF;^26–29^ therefore, we posited that VWF binding could account for the loss of free PS. VWF, in its shear-induced unfolded state, binds several blood proteins, including platelet glycoprotein Ib (GPIb)^30^ and ADAMTS13 (A Disintegrin And Metalloproteinase with ThromboSpondin type 1 motifs, member 13).^31^ To identify shear-dependent VWF-binding proteins, we incubated plasma samples with VWF-derivatized beads, with or without vortexing for 30min to mimic shear and maximize unfolding, and identified bound proteins by mass spectrometry. In all three plasma samples (two pooled, one single-donor), vortexing increased the quantity of PS bound to VWF (>10,000,000-fold in 2 of 3 samples), suggesting that shear-induced unfolding of VWF exposes a PS-binding site (**Figure 1A-B**). By comparison, albumin and fibrinogen only increased 2-4-fold after vortexing. C4BP-α was also enriched (**Figure 1C**), though to a lesser extent than PS (5-38-fold). However, C4BP-β, the PS-binding subunit, was not detected. This suggests that C4BP-bound PS does not bind VWF. Factor VIII was also not detected, likely because it remained bound to plasma VWF and did not transfer to the beads. There was a less than 2-fold increase in VWF-specific peptides, indicating that plasma VWF did not bind to the VWF-derivatized beads.

**Figure 1.**
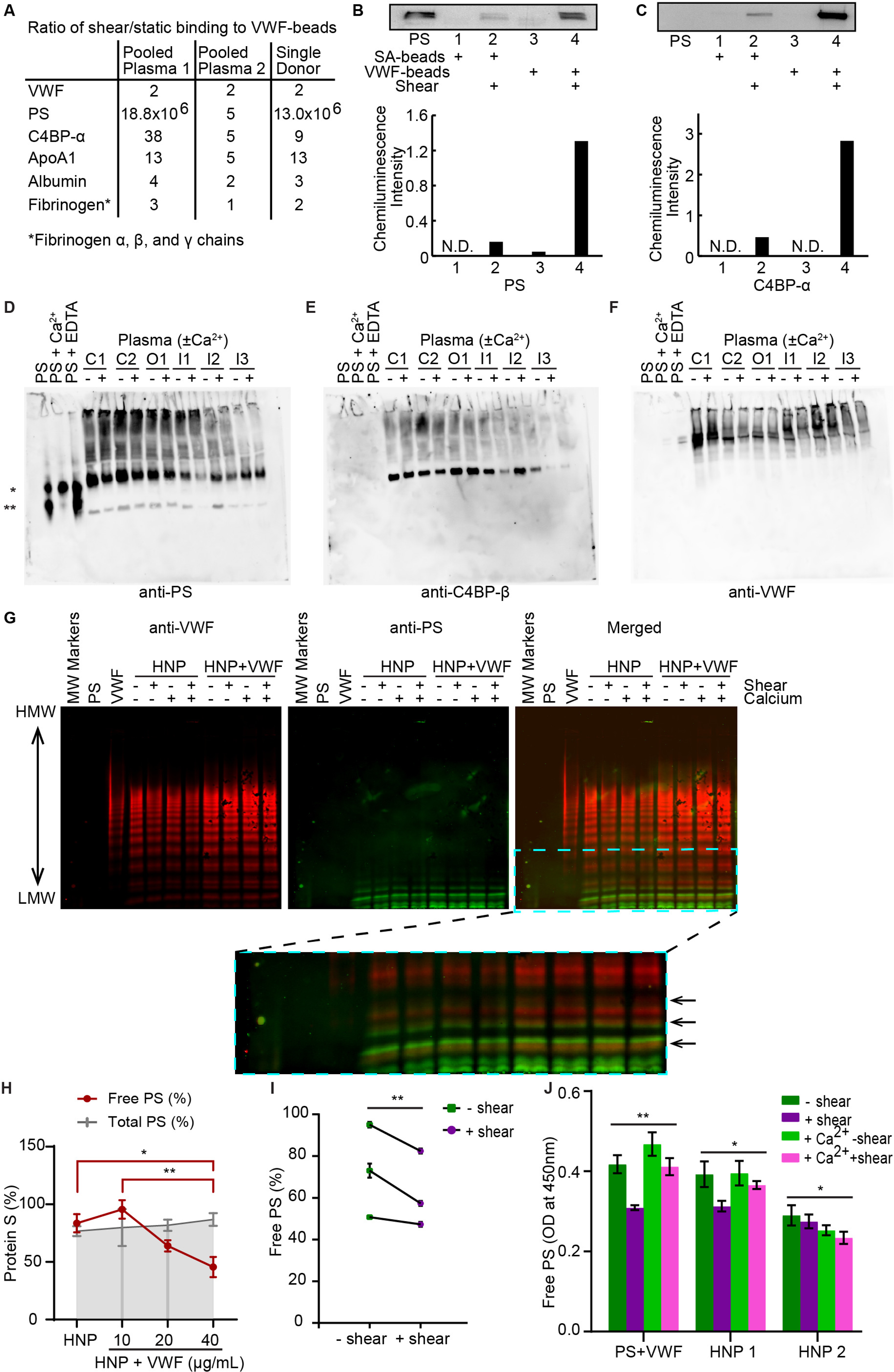
Sheared VWF interacts with PS in plasma and interferes with free PS measurement. (**A-C**) Streptavidin beads-immobilized biotinylated VWF was exposed to pooled human plasma or single donor plasma, under static conditions or under shear (vortexing), and bound proteins were analyzed with nanoLC-MS/MS, and probed for PS and C4BP-α by immunoblot, with serum albumin (SA) used as a control. (**A**) Mass spectrometry analysis of VWF pull-down in pooled normal plasmas (Pooled plasma 1 from Innovative Research, Pooled plasma 2 from Precision Biologics) and single donor. (**B**) Western blot of VWF pull-down probed for PS, and **(C)** reprobed for C4BP-α. (**D-F**) Purified PS (50ng), with either 20mM CaCl_2_ or 5mM EDTA, or 4μL citrated plasma (C1-2: healthy controls, O1: COVID-19 outpatient, I1-3: COVID-19 inpatients), with or without 1μM Hirudin, 5mM GPRP peptide, and 5mM CaCl_2_, were separated with native gel electrophoresis and probed or reprobed with (**D**) anti-PS, (**E**) anti-C4BP-β, and (**F**) anti-VWF antibody (*, apparent monomer band; **, apparent dimer band). (**G**) 25ng of PS or 10ng of VWF purified proteins, and 1μL of healthy normal plasma (HNP from Siemens), with or without shearing (∼2,500rpm for 30sec), or 1μM Hirudin, 5mM GPRP peptide, and 5mM CaCl_2_ supplementation, or additional 10μg/mL of purified VWF, were separated for VWF multimer analysis using SDS-agarose electrophoresis. The largest band on the molecular weight marker was 460kDa. VWF is shown on red channel, PS on green, and zoomed-in inset of the merged image is presented for clarity. (**H**) Free PS and total PS ELISA measurements of healthy normal plasma (HNP, from Siemens), with additions of purified VWF, under shear (∼2,500rpm for 30sec). (**I**) Free PS ELISA measurements of healthy control plasmas from single donors with or without shear. *P*-values are according to Wilcoxon matched-pairs signed rank test (**, *p*<0.01) **(J)** Free PS ELISA results of purified proteins (200nM PS and 10μg/mL VWF in HBSA), HNP1 (Corgenix), and HNP2 (Siemens), with or without shear or 1μM Hirudin, 5mM GPRP peptide, and 5mM CaCl_2_ supplementation. N.D.; non-detectible. *P*-values for (H) are according to two-way ANOVA with Sidak’s multiple comparisons test. *P*-values for (I-J) are according to Kruskal-Wallis with Dunn’s multiple comparison test. \**p*<0.05; \*\**p*<0.01

To detect PS binding partners, we analyzed plasma from healthy individuals and Coronavirus Disease 2019 (COVID-19) patients on native, non-denaturing gels and probed for PS-containing complexes by immunoblotting (**Figure 1D**). PS dimerizes in the absence of calcium, when at high concentrations (17µM); however, such concentrations are not reached in plasma.^32,33^ In **Figure 1D**, plasma PS migrated in a pattern distinct from purified PS (left 3 lanes), with no species co-migrating at the same apparent mobility as either the PS monomer (*) or dimer (**). Instead, several new complexes appeared, most above the monomer, and many toward the top of the gel. The most intensely stained band co-migrated with a band also detected by an anti-C4BP-β antibody (**Figure 1E**), consistent with reports that ∼60% of PS is complexed with C4BP.^4^ Other PS-positive bands co-migrated with bands detected by antibodies that recognize other known PS-binding proteins (e.g., Mer, PC, TFPI, factor V (**Figure S1**)) or VWF (**Figure 1F**), each of which migrated differently in plasma than with purified protein (**Figure S2**).

PS (∼69 kDa as a monomer) was also observed on VWF multimer gels (**Figure 1G, S4A**) and ran as a ladder of bands, similar to that seen for VWF. While the lowest molecular weight band migrated similarly to a complex with C4BP (**Figure S3B**), the remaining bands co-migrated with the low molecular weight VWF multimers (**Figure 1G, S3A**). This was apparent regardless of shearing or calcium supplementation, implying either a presence of pre-existing PS/VWF complexes in plasma, or that VWF unfolding in the electrophoretic buffer or system enabled PS binding. Overall, the results indicate that PS has multiple binding partners in plasma, apart from C4BP, including VWF.

### Soluble VWF interacts with PS in plasma and blocks free PS measurement

We next assessed whether sheared VWF affected free PS measurement in plasma. VWF was sheared by vortexing and added to plasma at different concentrations (0, 10, 20, and 40μg/mL) to mimic physiological (∼10μg/mL) and pathological (∼40μg/mL) concentrations. For these experiments, VWF was vortexed for a shorter time (30sec) to prevent loss of VWF and mimic milder unfolding conditions. In **Figure 1H**, VWF dose-dependently interfered with the detection of free, but not total, PS in pooled plasma. Similar vortexing of individual plasmas reduced free PS in all samples, though the extent of decrease varied between individuals (**Figure 1I**). The effect of vortexing was similar either in the absence or presence of calcium when measured with either purified proteins or plasma (**Figure 1J**). Vortexing did not alter the VWF concentration (**Figure S4**).

Next, PS and VWF were each derivatized with the photoactivatable, biotin transfer reagent Sulfo-SBED. The labeled proteins (PS-SBED and VWF-SBED) were incubated with PS-immunodepleted plasma, plasma from a type III von Willebrand Disease (VWD) patient, or pooled healthy donor plasma. Crosslinking was induced with exposure to UV light, biotinylated proteins were isolated with streptavidin-coated beads, and analyzed by immunoblotting (**Figure 2, S5**). As expected from the data in Figure 1, both proteins interacted with multiple binding partners in plasma (**Figure S5A**). PS-SBED transferred biotin to VWF, and as VWF-SBED did to PS (**Figure 2**). This transfer was increased upon vortexing, suggesting a direct shear-dependent interaction. Neither protein interacted non-specifically with the streptavidin beads (**Figure S5B**). Crosslinked complexes were too large to enter the gel under nonreducing conditions.

**Figure 2.**
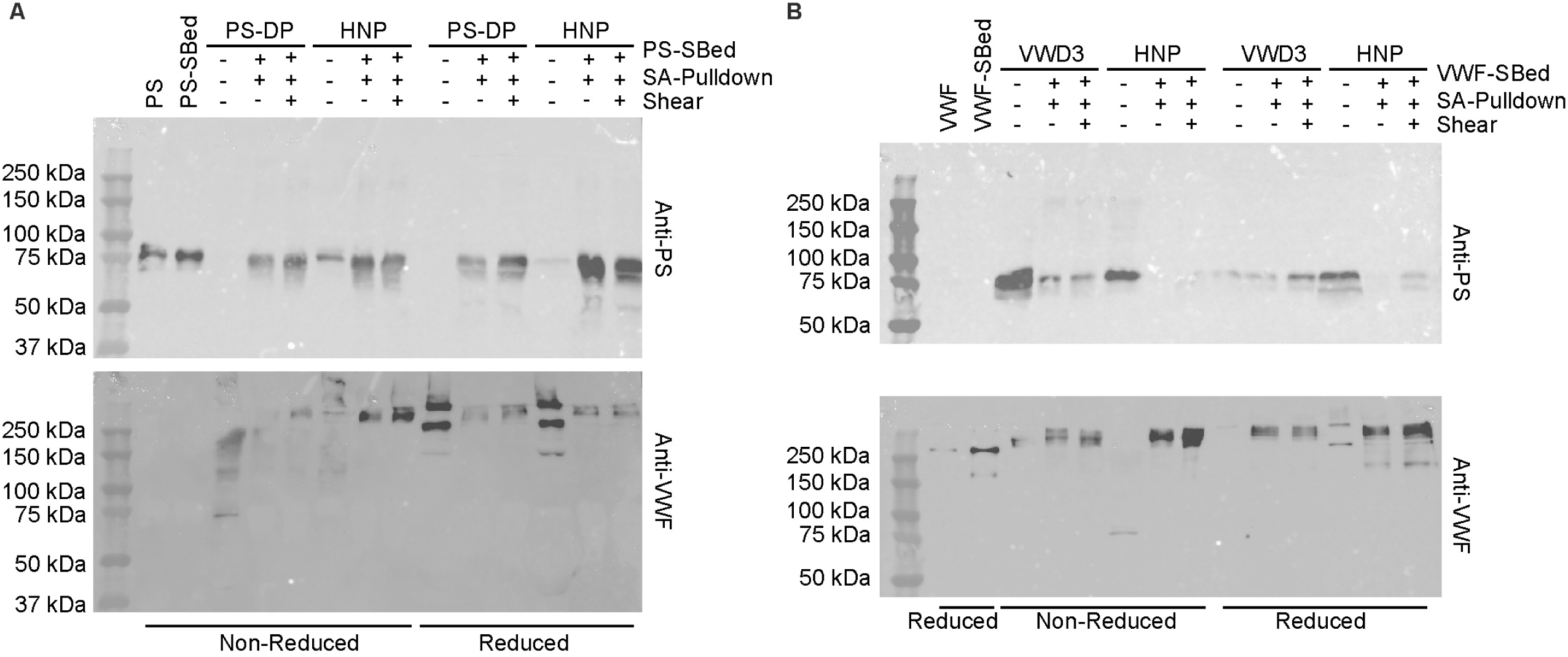
Labeled PS and VWF interact directly in human plasma in a shear-dependent manner. Sulfo-SBED-labeled PS (25µg/mL) (**A**) or VWF (10µg/mL) (**B**) was incubated in PS-depleted plasma (PS-DP) or plasma from a patient with Type III VWD (VWD3), respectively, or in pooled healthy normal plasma (HNP), in the presence or absence of shearing. Crosslinking was induced by exposure to UV light (365nm), and biotinylated proteins were isolated using streptavidin-coated agarose beads. Eluted proteins were identified by immunoblotting under non-reducing or reducing conditions, probed with antibodies against PS (top) or VWF (bottom).

### VWF competes with TFPI, but not APC for binding PS

To further define the PS/VWF interaction, we developed a PS/VWF complex ELISA, in which VWF is captured by a polyclonal antibody and bound PS is detected with a second polyclonal antibody. Using this assay, captured VWF bound PS from plasma in a time-dependent manner (**Figure 3A**), consistent with VWF immobilization inducing exposure of protein binding sites.^34^ Vortexing and calcium had no effect. Similarly, biotinylated PS bound to immobilized VWF, as detected with Streptavidin-HRP (**Figure 3B**). Pretreatment with a polyclonal anti-PS antibody reduced detection of the PS/VWF complex, as did the addition of saturating concentrations of TFPIα (**Figure 3B**). However, the addition of APC had no effect. These data suggest that VWF likely binds PS at or near the TFPIα binding site, which is located within the sex hormone-binding globulin (SHBG) region. APC binds within the PS epidermal growth factor (EGF) domains. Saturating concentrations of MerTK increased PS detection (**Figure 3B**). As MerTK preferentially binds dimerized PS,^35^ the increase may reflect the presence of PS dimer or indicate that dimeric PS preferentially binds VWF.

**Figure 3.**
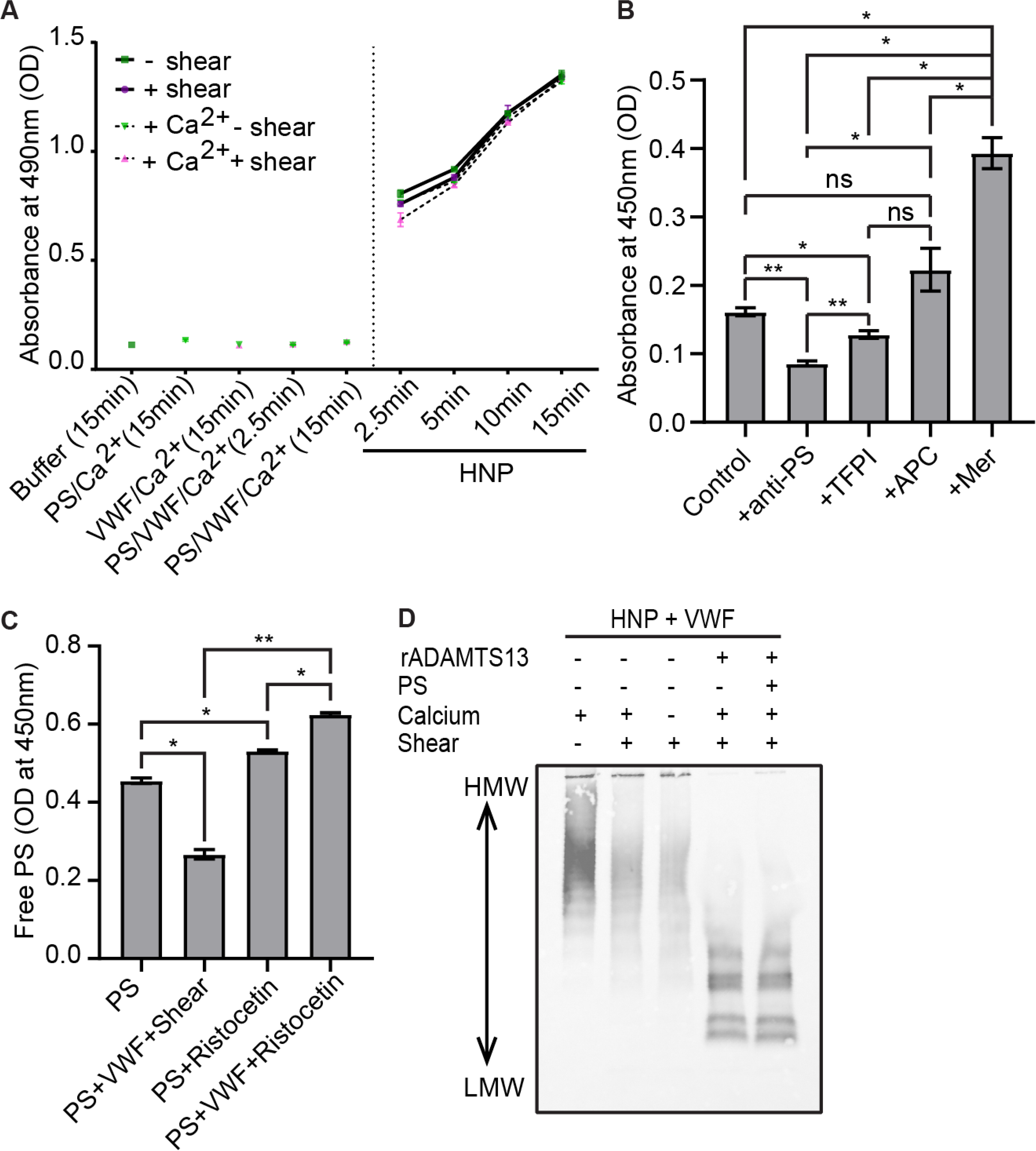
Direct measurement of the PS/VWF complex in plasma. (**A**) PS/VWF complex was measured by ELISA, using polyclonal antibodies to capture VWF and detect bound PS. Experiments were performed with purified proteins (200nM PS, 10μg/mL VWF, 5mM CaCl_2_, as indicated, in HBSA) or healthy normal plasma (HNP), with or without shear or 1μM Hirudin, 5mM GPRP peptide, and 5mM CaCl_2_ supplementation. Samples were incubated on the plate for the indicated time. (**B**) Biotinylated PS (150nM) was added to PS-immunodepleted plasma, and PS/VWF complex was detected as in (A), except using Streptavidin-HRP to detect. Experiments were performed in the presence or absence of saturating concentrations of an anti-PS polyclonal antibody, TFPIα, APC, or MerTK. (**C**) Free PS was measured, as in Figure 1, using purified PS (200nM), with or without purified VWF (10μg/mL) and Ristocetin (2mg/mL). (**D**) VWF multimer blot of 0.25μL HNP with additional 150nM purified VWF, with or without 50nM recombinant ADAMTS13, 200nM PS, or 1μM Hirudin, 5mM GPRP peptide, and 5mM CaCl_2,_ supplementation with or without shear (∼2,500rpm for 60min). Every data point is the average of three replicates (mean±SD). *P*-values for (**B**) and (**C**) are according to Kruskal-Wallis with Dunn’s multiple comparison test (\**p*<0.05; \*\**p*<0.01).

The PS interaction site on VWF appears distinct from that which binds GPIb. Ristocetin, which exposes the GPIb binding site had no effect (**Figure 3C**). In addition, exogenous PS did not interfere with VWF cleavage by ADAMTS13 (**Figure 3D**), which targets the unfolded A2 domain. Collectively, the data indicate that PS binds a shear-exposed site on VWF that is distinct from those that bind GPIb and ADAMTS13.

### PS/VWF complex is stable under arterial flow conditions

Microfluidics experiments were performed to visualize PS and VWF on thrombi formed under shear. Whole blood was supplemented with fluorescently tagged PS and VWF, either with or without prior shearing, recalcified, and perfused at 35dyn/cm^2^ (∼arterial) shear stress (**Figure 4A-B**). In the absence of prior shearing, no apparent colocalization of PS and VWF was observed, as PS and VWF bound independently to different thrombi structures (r=0.177).

**Figure 4.**
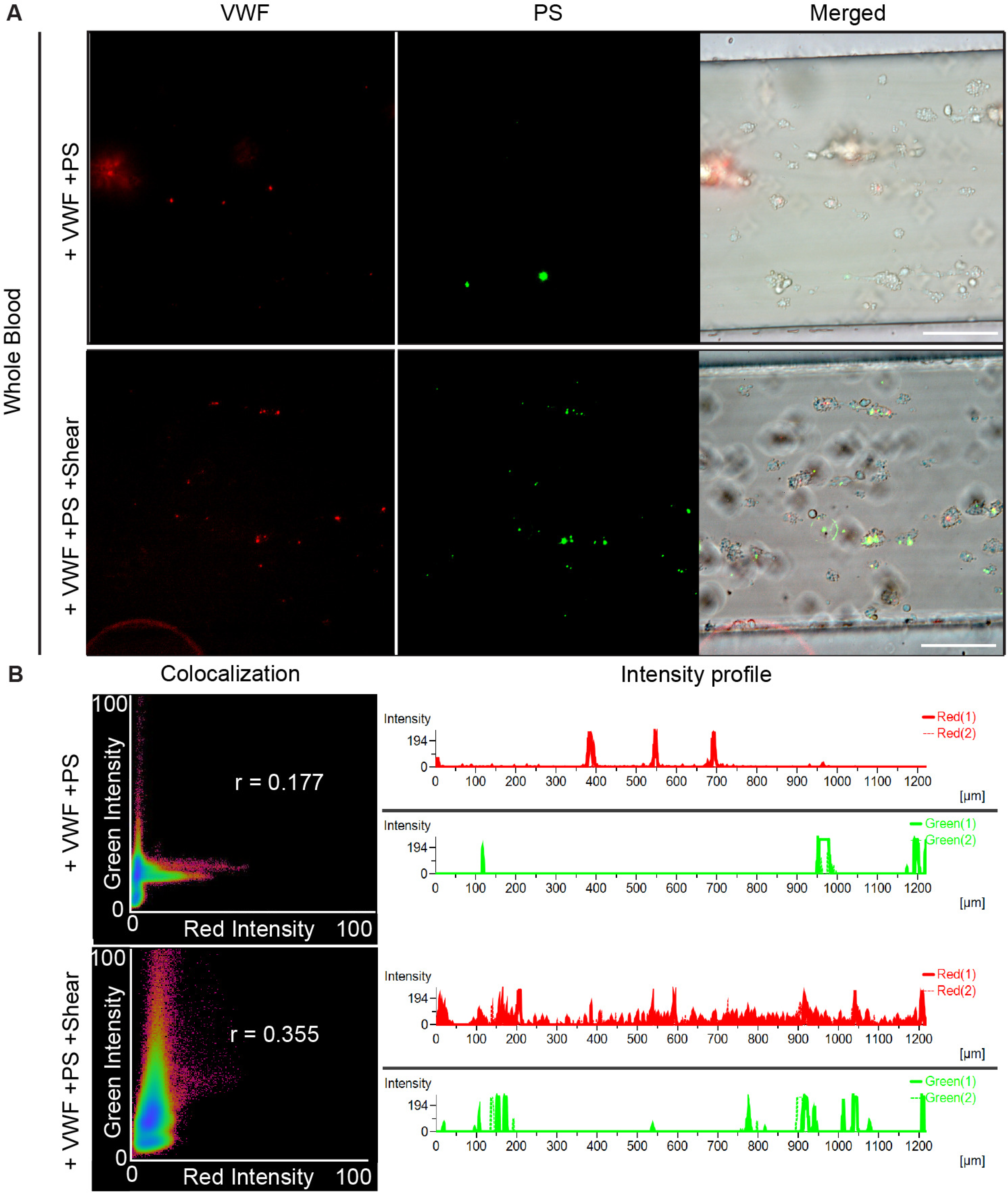
The PS/VWF complex is stable under flow. Whole blood was supplemented with 10μg/mL AlexaFluor555-conjugated VWF and 200nM AlexaFluor488-conjugated PS and perfused through collagen-coated channels at 35dyn/cm^2^. Experiments were performed either without or with PS/VWF vortexing. (**A**) Images taken after flushing with buffer. Scale bar=100 μm. (**B**) Colocalization (shown are Pearson correlation r) and intensity profile analyses using the NIS Element software.

However, when PS and VWF were sheared to form the complex before addition, the two proteins co-localized on platelet thrombi (r=0.355), indicating that the complex is stable under flow. However, arterial laminar flow did not induce complex formation alone.

### PS/VWF complex forms under turbulent flow, in a calcium-sensitive manner

To investigate the effect of turbulent or disrupted flow on PS/VWF binding, we perfused VWF through a PDMS microfluidic device^36^ that allows visualization of VWF self-association around the PDMS block (**Figure 5A**). VWF and fluorescently labeled PS were perfused into the microfluidic device in the absence (**Figure 5B, S6A, Videos S1-2**) or presence (**Figure 5C, S6B, Videos S1-2**) of 2mM calcium chloride. Both PS and VWF have calcium-sensitive features;^37–39^ thus, we assessed the effects of calcium on the PS/VWF interaction using purified proteins. We observed PS binding to VWF under shear and found that PS binding was significantly higher in the absence of calcium (**Figure 5C**). As the A2 domain unfolds more readily in the absence of calcium, this suggests the PS-binding site is within A2.^37,39^ As expected, we observed no fluorescence intensity over background in the absence of VWF or PS (**Figure 5C, S6C, Videos S1-2**). To account for differences in VWF self-association with and without calcium, we normalized PS binding to the area of VWF self-association in each channel. Even with normalization, PS binding was significantly higher in the absence of calcium (**Figure 5D**).

**Figure 5.**
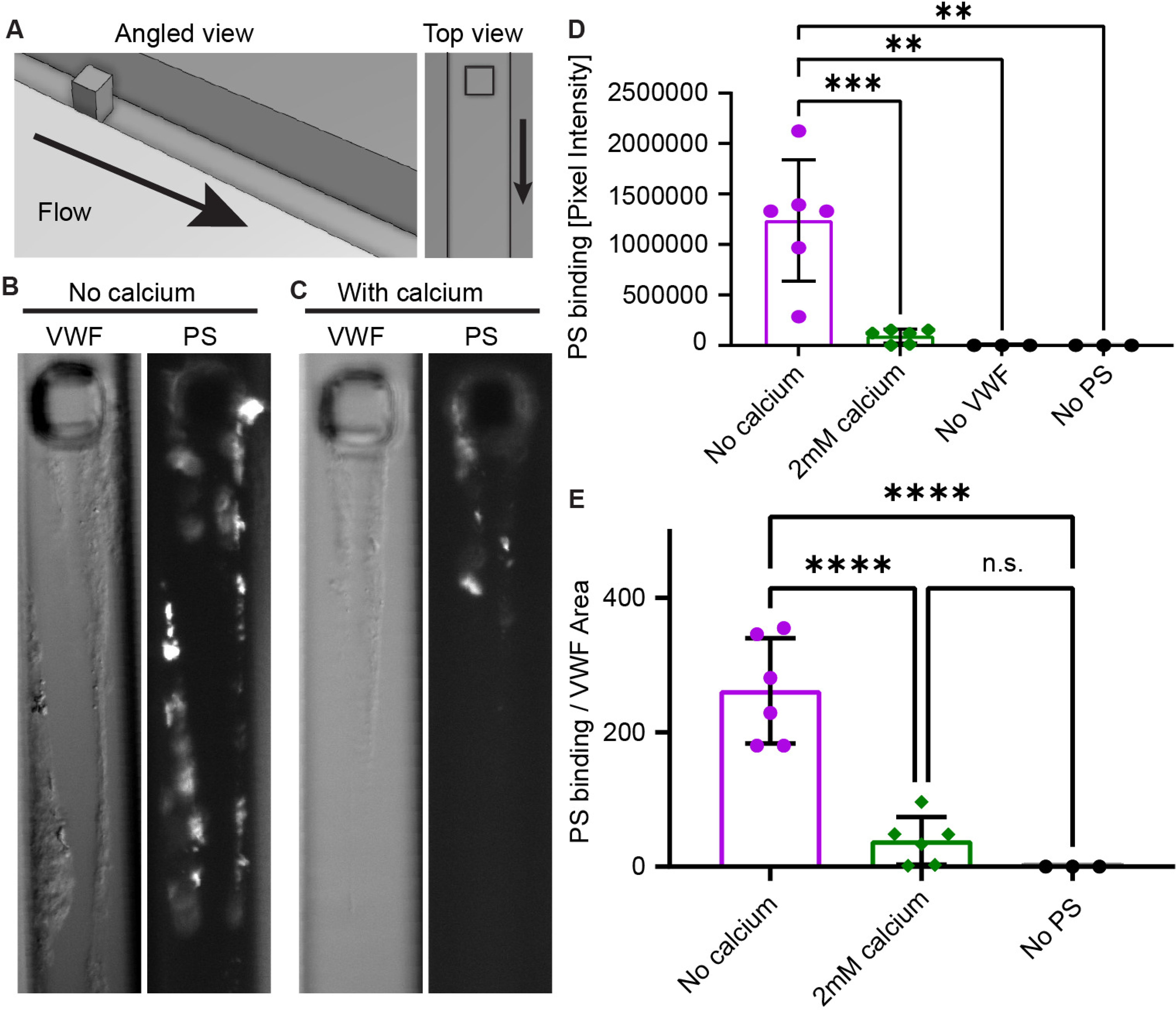
PS binds to VWF that unfolds during disrupted flow conditions, in a calcium-modulated manner. VWF and PS were perfused through a PDMS microfluidic device (**A**). In the absence (**B**) or presence of 2mM calcium (**C**), VWF self-association and Alexa Fluor 488-PS binding were visualized by DIC and epifluorescence microscopy, respectively. (**D**) Raw pixel intensity above noise was summed for these conditions and controls without VWF (with PS) and without PS (with VWF). Each data point indicates an independent run. (**E**) When accounting for the area of VWF self-association, PS binding was significantly higher than all other conditions. Error bars are SD. Statistical significance was determined with an ANOVA and Tukey post hoc test (n.s., non-significant; **, *p*<0.01; ***, *p*<0.001; ****, *p*<0.0001). Scale bar=25μm.

### Unfolded VWF reduces PS anticoagulant activity

The effect of VWF unfolding on PS anticoagulant activity was measured using plasma thrombin generation in the presence or absence of exogenous APC (5nM), as described by Brugge *et al.*,^40^ and with or without vortexing to generate shear (**Figure 6A**). The data were expressed as percent reduction in peak thrombin upon APC addition. In pooled plasma, APC reduced peak thrombin by 72.0±7.1%. By contrast, a 3.0±3.5% reduction was observed in PS-immunodepleted plasma. Vortexing plasma decreased the apparent anticoagulant activity, as APC only reduced peak thrombin by 56.0±7.8% (p=0.005). Similar results were observed in plasma from an individual donor. This reduction was entirely dependent on PS, as APC had no effect in vortexed PS-immunodepleted plasma (1.7±1.0% reduction, p=0.497). The effect was also dependent on VWF. In Type III VWD plasma, PS had increased apparent anticoagulant activity (reducing the peak by 87.2±3.3%), which was not changed by vortexing (83.2±0.5%).

**Figure 6.**
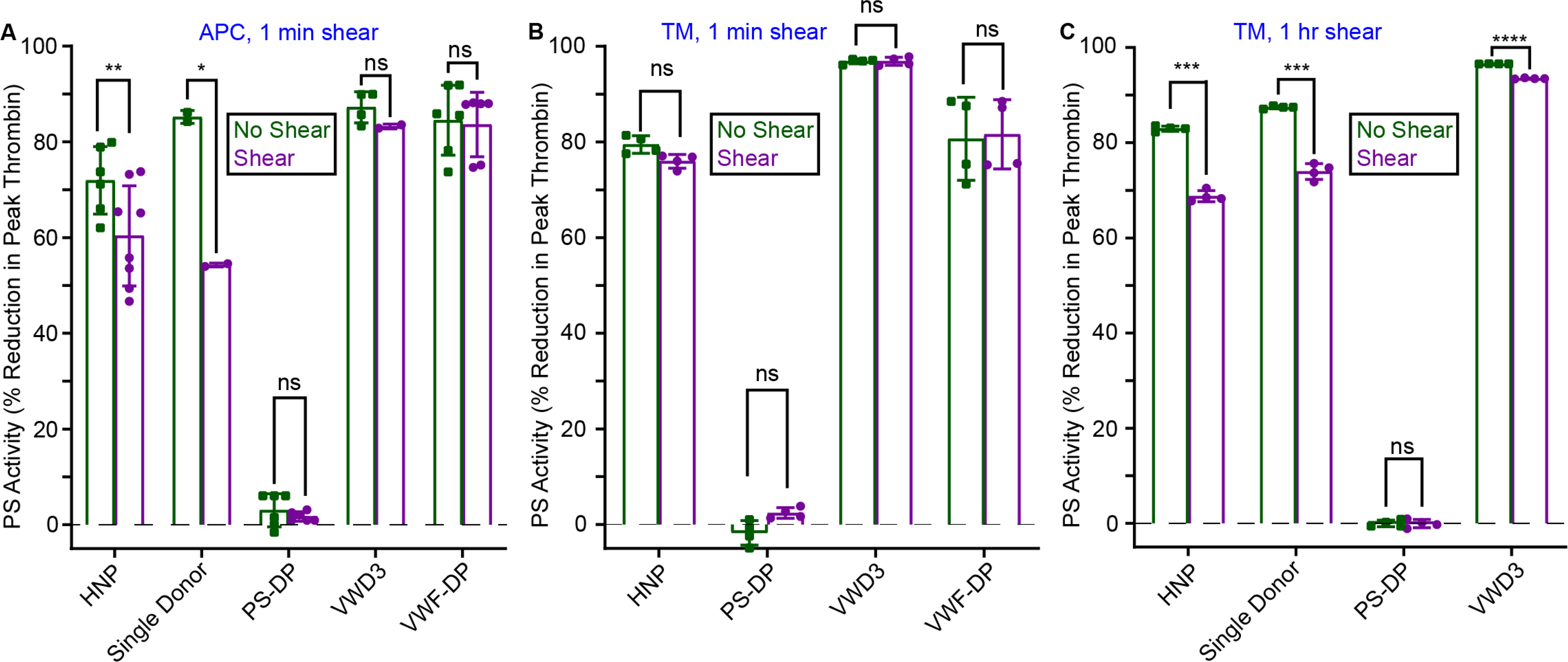
Unfolded VWF reduces PS plasma anticoagulant activity. Plasma thrombin generation was measured in pooled healthy normal plasma (HNP), single donor plasma, PS-depleted plasma (PS-DP), plasma from a patient with Type III VWD (VWD3), or VWF-depleted plasma (VWF-DP) in the presence or absence of exogenous APC (5nM, **A**) or thrombomodulin (20nM, **B-C**) and the presence or absence of short-term shear (1min, **A-B**) or long-term shear (1hr, **C**). Paired t-tests were performed with each individual data set, comparing plasma with or without shearing (\**p*<0.05, \*\**p*<0.01, \*\*\**p*<0.001, \*\*\*\**p*<0.001).

Next, we utilized a recently described PS-sensitive thrombin generation assay that relies on *in situ* protein C activation (**Figure 6B-C**).^11^ In contrast to the APC experiments, vortexing plasma for 1min had little effect on anticoagulant activity. However, vortexing for 1hr, to maximize VWF unfolding, resulted in decreased APC/PS activity. As with the exogenous APC experiments, this reduction was dependent on PS and VWF. Collectively, the *in vitro* data indicate that unfolded VWF binds PS, blocks the measurement of free PS, and reduces PS anticoagulant activity.

### Free PS Deficiency in COVID-19 Patients Correlates with Increased VWF and Thrombin Generation Potential

To assess whether this interaction occurs *in vivo*, we analyzed plasma samples from a cohort of healthy controls (n=38) and patients with severe COVID-19 (ICU patients, n=30), a condition associated with free PS deficiency.^18^ As expected, samples from COVID-19 patients exhibited elevated signs of infection and inflammation compared to controls, including elevated anti-spike protein IgG (Endpoint Titer=18,500±23,200 in patients compared to 199±361 in controls, expressed in reciprocal dilution; *P*<0.0001; **Figure S7A**), tumor necrosis factor (1.33±1.01 *vs.* 0.78±0.87pg/mg total protein, *P*=0.003; **Figure S7B**), interleukin-6 (2.56±5.18 *vs.* 0.043±0.038pg/mg total protein, *P*<0.0001; **Figure S7C**), activated complement component C5 (1,195±1,021 *vs.* 179.8±174.6pg/mg total protein, *P*<0.0001; **Figure S7D**), and plasma myeloperoxidase (15.98±9.19 *vs.* 2.72±1.15pg/mg total protein, *P*<0.0001; **Figure S7E**). Anti-spike IgG was detected in some controls who had received the first dose of a COVID-19 vaccine. Markers of coagulopathy were also detected in the patients, including elevated plasma TF activity (2.55±4.15 *vs.* 0.76±0.7fmol/g total protein, *P*=0.005; **Figure S7F**), monocyte TF expression (2,360±1,669 *vs.* 1,259±403MFI, *P*<0.0001; **Figure S7G**), plasma D-dimer (72.67±141.7 *vs.* 5.18±4.39ng FEU/mg total protein, *P*<0.0001; **Figure S7H**), and plasma microclots, detected by Thioflavin T staining^41^ (0.072±0.092 *vs* 0.03±0.051, *P*<0.0001; **Figure S7I**). All of these were consistent with the hypercoagulable and hyperinflammatory state found in COVID-19 patients.^42–45^

Total PS was unchanged (*P*=0.611) (**Figure 7A**), but free PS (**Figure 7B**) decreased in patients (81±40.6%) compared to controls (100±33%) (*P*=0.017). The free PS deficiency could not be explained by C4BP, as C4BP-β (**Figure 7C**) was unchanged in patients, nor by PC (**Figure S8A**). Soluble Mer-TK was elevated in patients compared to controls (135.8±80.1 *vs.* 77.3±45pg/mg total protein, *P*=0.0002; **Figure S8B**), but this increase was not sufficient to explain the decreased free PS (picomolar concentrations of Mer-TK (70-1000pM in controls and patients) *vs.* nanomolar free PS (5-15μg/mL or ∼150nM)).

**Figure 7.**
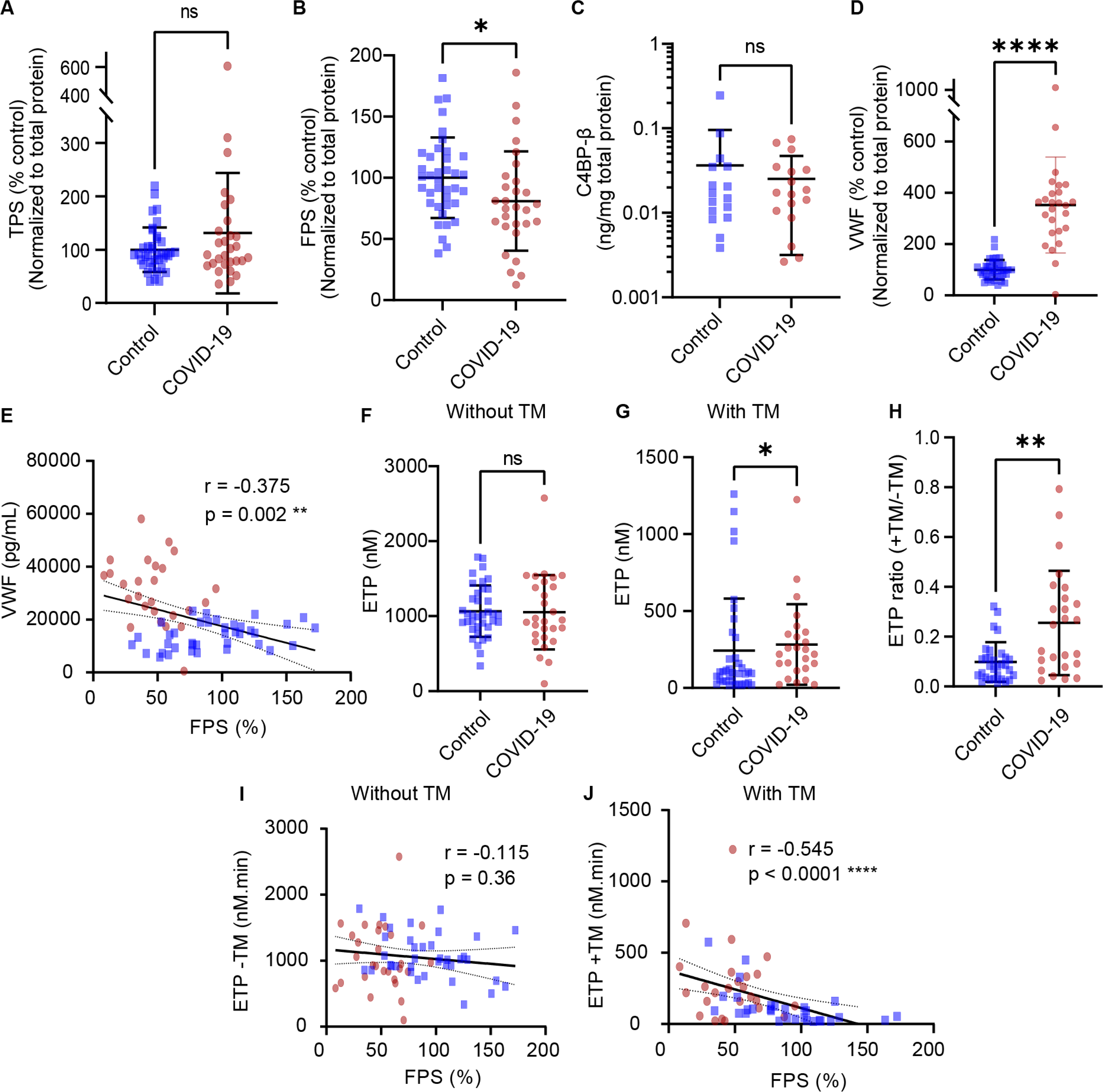
Free protein S (PS) deficiency in COVID-19 patients can be explained by changes in VWF but not C4BP. Citrated plasma samples were collected from healthy controls (n=38) and COVID-19 patients (n=30). Due to sample limitations, some measurements could not be performed with all subjects. (**A**) Total PS, (**B**) free PS, (**C**) C4BP-β, and (**D**) VWF were measured by ELISA. (**E**) Free PS plotted against VWF:Ag. (**F-J**) Plasma thrombin generation was measured using calibrated automated thrombography. Several patients were receiving heparin prophylactic dose at the time of blood collection. Shown are the endogenous thrombin potentials (ETP) in assays initiated with 4μM phospholipids and 1pM tissue factor (TF) (**F**) or with 20nM thrombomodulin (TM) supplementation to evaluate the contribution of APC/PS activity (**G**), and (**H**) the ETP ratio (value with TM/value without TM). (**I-J**) Free PS plotted against ETP in the absence (**I**) or presence (**J**) of thrombomodulin. Every data point is the average of three replicates (mean±SD). (**A-H**) *P*-values are according to Mann-Whitney test (*, *p*<0.05; **, *p*<0.01). (**I-J**) *P*-values and r correlation coefficients are according to Spearman correlation analysis (***, *p*<0.001; ****, *p*<0.0001). Blue squares represent controls and red circles represent COVID-19 patients.

Plasma VWF antigen concentration (VWF:Ag) was elevated in the patients (353±187%, *P<*0.0001) compared to controls (100±38.2%) (**Figure 7D**), as previously reported.^41^ Soluble E-selectin was also elevated (191±72.5 vs 133±32pg/mg total protein in patients and controls, *P=*0.0001), consistent with increased VWF secretion from activated endothelium (**Figure S9A**). VWF multimer distribution was unchanged (**Figure S9B**). VWF:Ag negatively correlated with free PS concentration (r=-0.375, *P*=0.002) (**Figure 7E**).

Plasma thrombin generation was measured in the presence or absence of thrombomodulin. Despite most patients receiving heparin prophylaxis at the time of sample collection, plasma thrombin generation was significantly elevated when measured in the presence of thrombomodulin (peak thrombin, *P*=0.01; endogenous thrombin potential [ETP], *P=*0.043) but not in its absence (**Figure 7F-G, S10**). There was no difference between groups in the absence of exogenous TF (**Figure S10**). ETP ratio, defined as the value with thrombomodulin divided by without thrombomodulin, was also increased in patients compared to controls (peak thrombin ratio; *P*=0.014; ETP ratio; *P*=0.001) (**Figure 7H**). These data indicate reduced APC/PS pathway activity in hospitalized COVID-19 patients. Consistent with our previous report,^11^ free PS only weakly correlated with peak thrombin and did not correlate with ETP in the absence of thrombomodulin (**Figure 7I, S10E**), but showed a strong negative correlation in the presence of thrombomodulin (peak thrombin, r=-0.475, *P*=0.0002; ETP, r=-0.545, *P*<0.0001) (**Figure 7J, S10F**), indicating a significant functional consequence of the free PS deficiency.

## Discussion

Free PS deficiency is a common complication of severe viral infections, including HIV-1, dengue, varicella, and COVID-19.^10–17,19^ The specific decrease in free PS indicates a concurrent increase in a PS-binding protein, with C4BP being the best-known candidate. The major isoform (∼80%) of C4BP contains 7α and 1β chains, with the other isoforms containing either 6α and 1β or exclusively 7α or 6α.^46^ The β chain binds PS.^47,48^ While C4BP is commonly increased during infection and inflammation, this change is mostly associated with elevated α chain, suggesting that the acute phase C4BP is unlikely to bind PS.^21^ Thus, we hypothesized that another plasma protein is altered during inflammation, binds PS, reduces “free PS” measurement, and reduces PS anticoagulant activity. Our attention focused on VWF, since VWF is elevated in disease states that often present with PS deficiency.^26–29^ VWF also precipitates in PEG,^49^ which has been historically used to separate the free and bound pools of PS.^20^ Here, we showed that VWF binds PS in a shear-dependent manner, blocks free PS measurement, and reduces PS anticoagulant activity. Further, PS binds VWF as it unfolds dynamically under flow and forms a stable complex that associates with platelet thrombi. Finally, PS deficiency in a cohort of COVID-19 patients can be explained by changes in VWF but not C4BP. Based on these data, we propose that dysfunctional VWF directly sequesters PS, promoting thrombin generation through the reduction of available anticoagulant activity. This is a heretofore unrecognized procoagulant activity of VWF, and to our knowledge the first description of anticoagulant regulation by VWF.

To understand the potential causes of free PS deficiency and test for other PS-binding proteins, we began by visualizing PS in plasma from COVID-19 patients and healthy controls by non-denaturing gel electrophoresis and immunoblotting (**Figure 1D-F**). A multitude of bands toward the top of the gel were observed in all plasma samples, indicating that PS circulates in plasma bound to many other proteins. Most of these interactions are likely lower affinity than C4BP and thus were not identified in previous studies using size exclusion chromatography.^4^ Mass spectrometry, immunoprecipitation, and Sulfo-SBED crosslinking analyses revealed increased binding of PS to VWF upon exposure to shear induced by vortexing the samples (**Figure 1A-C, 2**), and indeed some of the PS bands on the native gel co-migrated with VWF. Sheared VWF dose-dependently decreased the measurement of free PS, as did shearing plasma (**Figure 1H-J**). VWF is an abundant, large multimeric protein with protein binding sites that can be exposed under high shear.^30^ VWF is also a prothrombotic acute phase reactant during inflammation, including in COVID-19 (**Figure 7D**).^50,51^ We hypothesized that shear-induced unfolding of VWF exposes a PS-binding site, as it does for platelet GPIb^30^ or ADAMTS13.^31^

Our data suggest that the VWF binding site on PS is likely adjacent to or sterically hindered by the C4BP-binding site, which is found within the C-terminal SHBG region, as this is the epitope recognized by the monoclonal antibody used to measure free PS,^52^ and C4BP-β chain was not detected by either mass spectrometry or immunoblotting (**Figure 1**). Consistent with this, saturating concentrations of TFPIα, which also binds the SHBG region, competed with VWF for binding PS (**Figure 3B**). APC, which binds the N-terminal EGF domains,^53–56^ had no effect. Interestingly, MerTK, which binds dimeric PS, appears to promote the PS/VWF interaction, suggesting that VWF may have a higher affinity for dimeric PS. The exact PS binding site on VWF is unknown; however, it is distinct from the GPIb binding site, as treatment with Ristocetin, which exposes that site, did not reduce free PS (**Figure 3C**). Finally, calcium disrupts the PS/VWF interaction (**Figure 5**), an observation with two evident interpretations: First, calcium stabilizes the folded state of A2^37,39^, and thus PS may bind unfolded A2; Second, calcium disrupts PS dimerization. Either or both mechanisms are consistent with our data. An interaction with A2 would also be consistent with the presence of PS on VWF multimer gels (**Figure 1G**), as PS may interact with terminal A2 fragments, exposed in smaller VWF multimers generated through cleavage by ADAMTS13.

Pre-formed PS/VWF complexes were stable in recalcified whole blood, and co-localized on platelet thrombi under arterial laminar flow (**Figure 4**), though these conditions did not induce formation of the complex *in situ*. In contrast, PS bound to VWF that unfolded and self-associated under turbulent shear flow conditions (**Figure 5**), such as hypothesized to occur in COVID-19 vasculopathy.^57^ VWF interfered with PS anticoagulant activity, as measured in two different plasma thrombin generation assays (**Figure 6**). Vortexing plasma resulted in a consistent ∼10% decrease in the anticoagulant effect of exogenous APC, an effect dependent on PS and VWF. Similarly, extended vortexing of plasma reduced the anticoagulant effect of exogenous thrombomodulin, an activity that was again dependent on PS and VWF. The thrombomodulin results indicate a time-dependence to the effect, in which VWF must be unfolded and able to bind PS at the time that APC is available. Under short-term shear conditions, VWF likely refolds during the 10min incubation time of the thrombin generation assay. Upon exposure to prolonged shear, as likely occurs in patients with chronic inflammation, sufficient VWF remains unfolded and/or bound to PS to reduce anticoagulant activity. To our knowledge, this is the first evidence of a direct effect of VWF regulating plasma anticoagulant function.

Our data offer an explanation for free PS deficiency during infections. Free PS deficiency was prevalent in our COVID-19 population (**Figure 7A-B**). However, it could not be explained by alterations in C4BPβ, which was unchanged in COVID-19 patients, as was PC (**Figure 7C, S8A**). By contrast, VWF was elevated in COVID-19 patients and inversely correlated with free PS (**Figure 7D-E**). The free PS deficiency also correlated with elevated plasma thrombin generation (**Figure 7F-J**). Despite prophylactic heparin anticoagulation received by some patients, plasma thrombin generation was not decreased in patients in the absence of thrombomodulin. In the presence of thrombomodulin, peak thrombin and endogenous thrombin potential were elevated in patients, signifying lower sensitivity to thrombomodulin and reduced APC/PS pathway activity.

As shearing reduces free PS in healthy control plasma, the effect of VWF on free PS is likely mediated by shear-dependent unfolding, rather than changes in VWF antigen. Unfolding and self-association of VWF are favored in regions of flow acceleration, as occur in stenotic arteries or valves and in resistance vessels,^58^ especially during hypertension. It is noteworthy that in two studies published early in the COVID-19 pandemic, hypertension was the most common comorbidity associated with severe disease,^59,60^ and that hypertension is associated with reduced PS.^61^

In conclusion, we propose a model in which shear-induced unfolding exposes a binding site on VWF, possibly within the A2 domain, that directly interacts with PS, likely within the SHBG domain. VWF binding reduces PS anticoagulant activity and blocks the measurement of free PS. This proposed mechanism is consistent with the free PS deficiency that develops in COVID-19, and with historic measurements of free PS deficiency, which is prevalent in inflammatory conditions that are also associated with VWF dysfunction, and which relied on PEG precipitation of plasma, a process also used to precipitate VWF. We anticipate that this mechanism broadly contributes to acquired free PS deficiency in inflammatory conditions in which VWF is exposed to elevated vascular shear force, and that VWF binding may modulate PS anticoagulant activity under normal hemostatic conditions when VWF unfolds at the site of vascular injury.

## Acknowledgments

This study was supported by NHLBI grants HL56652, HL138179, and HL150818 (S.W.W.), HL129193 (J.P.W.), HL145262 (J.A.L), HL007093 (M.Y.M.), VA Merit I01BX003877 (S.W.W.), and UK CCTS pilot grants (J.G.W. and J.P.W.). The UK Flow Cytometry & Immune Monitoring core facility is supported in part by the Office of the Vice President for Research, the Markey Cancer Center, and an NCI Center Core Support Grant (P30 CA177558). This study was supported by the National Center for Research Resources and the National Center for Advancing Translational Sciences, National Institutes of Health, through Grant UL1TR001998. We thank Dr. Etheresia Pretorius for her advice about the Thioflavin T procedure.

## Author Contributions

M.M.S.S. and J.P.W. designed and performed experiments, analyzed data, and wrote the first draft of the manuscript. M.Y.M., H.R.A., M.H., D.W.C., X.Y.F., S.G., D.M.C., K.S.P., M.B., C.P., and X.L. designed and performed experiments and analyzed data. J.P.W., J.A.L., J.G.W., B.A.G., and S.W.W. designed the study and critically revised the manuscript. D.W.C., D.F.D., and Z.Z. critically revised the manuscript. A.C.T., J.Z.P., J.L.S., G.A.S., M.B.B., and K.S.C. recruited the patients. All authors reviewed the final version of the manuscript and approved its submission.

## Supplemental Materials

Supplemental Methods

Figures S1-S10

Videos S1-S2

---

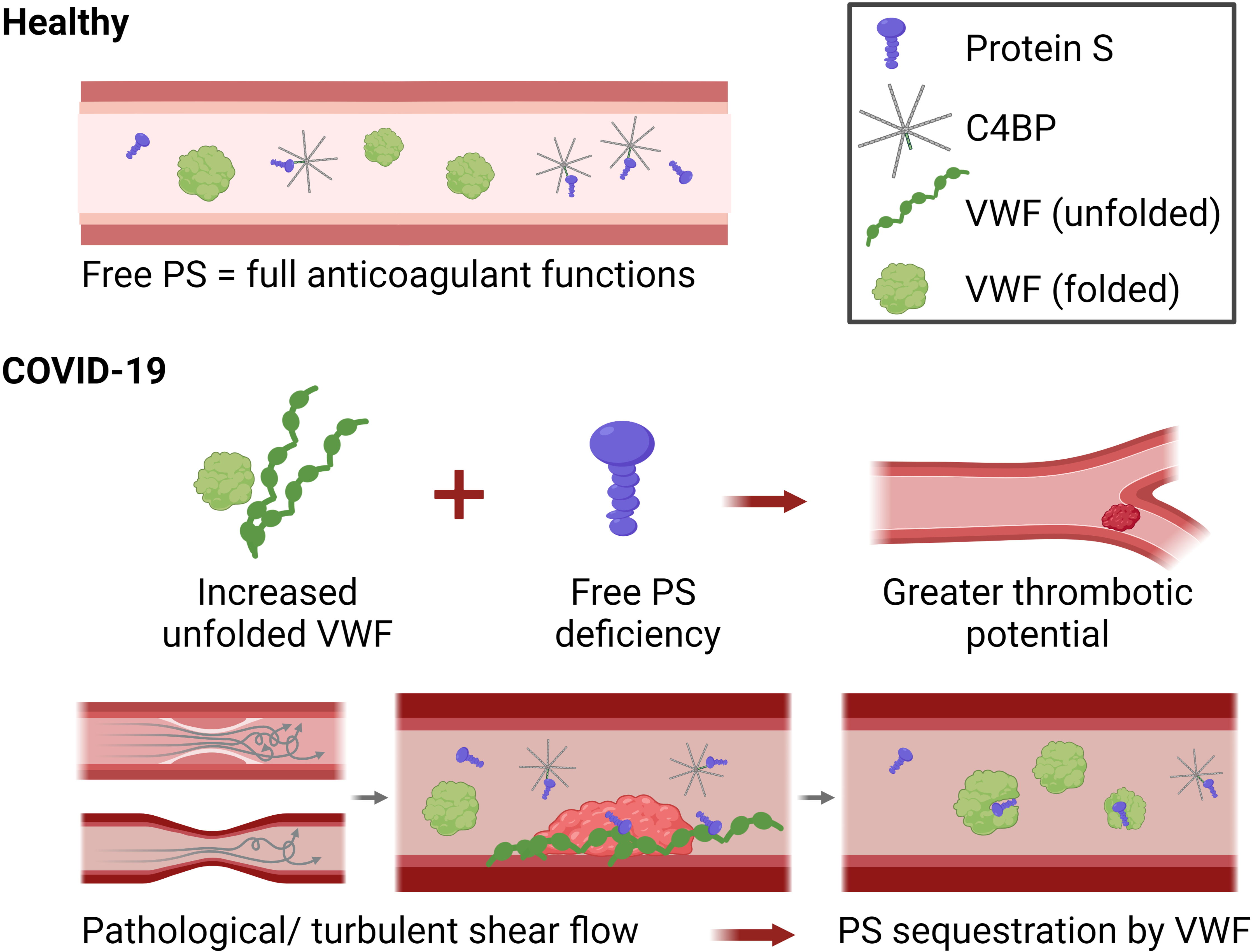

