## Supplemental Methods and Figures S1-S10 for "Unfolded Von Willebrand Factor Binds Protein S and Reduces Anticoagulant Activity"

### Materials.

Pooled healthy normal plasma (HNP; control plasma N) and human tissue factor (TF) (Dade Innovin) were from Siemens Healthineers (Erlangen, Germany). Thioflavin T was from Cayman Chemical (Ann Arbor, Michigan, USA). Human plasma-purified PS, FVIIa and factor X (FX) were from Enzyme Research Laboratories (South Bend, IN, USA). Human VWF (factor VIII free) was from Haematologic Technologies, (Essex Junction, VT, USA). Fibrinogen-, PS-, and VWF-immunodepleted plasma were from Affinity Biologicals (Ontario, Canada). Von Willebrand Disease type 3 deficient plasma was from HRF, Inc (Raleigh, NC, USA). Human thrombomodulin was from PeproTech (Cranbury, NJ, USA). Phosphatidylcholine (PCho), phosphatidylethanolamine (PEth), and phosphatidylserine (PSer) were from Avanti Polar Lipids (Alabaster, AL, USA) and were used to prepare PCho:PEth:PSer (40:40:20) vesicles (= phospholipids/ PL) according to the protocol of Morrissey.[1] Thrombin calibrator and FluCa substrate for Calibrated Automated Thrombography (CAT) were from Diagnostica Stago (Asnières-sur-Seine, France). O-phenylenediamine dihydrochloride (OPD) substrate was from Sigma Aldrich (Ann Arbor, Michigan, USA), and Spectrozyme FXa from BioMedica Diagnostics (Windsor, Canada).

### Plasma pull-down of VWF-binding proteins

Recombinant biotinylated VWF, expressed by stably transfected HEK293 cells, was immobilized onto streptavidin magnetic beads (New England Biolabs).[22] The VWF-bound beads were exposed to pooled or single-donor human plasma, under static conditions or shear (vortexing at 37˚C for 30min) in the presence of 10mM EDTA. The beads were washed five times with PBS and then subjected to on-bead trypsin digestion and the released peptides were analyzed by nano-liquid chromatography coupled with tandem mass spectrometry (nanoLC-MS/MS), as described.[23] Proteins were identified by the presence of at least two tryptic peptides and the average peak area of the three most abundant unique peptides was used to calculate the abundance of the captured protein, expressed as the shear-to-static ratio. Bead-bound proteins were electrophoresed on reducing 4-20% SDS-PAGE gels, transferred to nitrocellulose, and probed with antibodies against PS, C4BP-β, and C4BP-α (LifeSpan Biosciences, Seattle, WA, USA).

**ELISAs**

ELISAs were used to measure human total PS (Enzyme Research Laboratories, IN, USA), free PS (Corgenix, CO, USA), protein C (Enzyme Research Laboratories, IN, USA), C4BP-β (LifeSpan Biosciences, WA, USA), soluble Mer (R&D Systems, MN, USA), D-dimer (Diagnostica Stago, Asnières-sur-Seine, France), Myeloperoxidase (R&D Systems, MN, USA), soluble E-selectin (R&D Systems, MN, USA), and VWF (R&D Systems, MN, USA), following manufacturers' instructions. Plasma IL-6 and TNF were measured using a Milliplex Human Cytokine Kit (Sigma Aldrich, St. Louis, MO, USA). Total PS and PC measurements were compared to standards of control plasma N, whereas free PS, C4BP-β, soluble Mer, D-dimer, MPO, soluble E-selectin, and VWF measurements used standards provided with the kits. Total PS ELISA utilizes paired polyclonal antibodies to PS, whereas free PS utilizes a monoclonal capture antibody that recognizes non-C4BP-β-bound PS. For some experiments, purified VWF was mixed with purified PS, or into control plasma N, with or without vortex-induced shearing (~2,500 rpm at RT for 30sec). For some experiments, 5mM $CaCl_2$ was added to plasma samples after 1mM hirudin (Sigma Aldrich, St. Louis, MO, USA) and 5mM gly-pro-arg-pro (GenScript Biotech, Piscataway, NJ, USA) peptide. For some experiments, 2mg/mL Ristocetin A sulfate

(MP Biomedicals, Irvine, CA, USA) was mixed with plasma. To correct for plasma dilution, measurements were normalized to total protein, determined by bicinchoninic acid assay (bioWORLD, Dublin, OH, USA).

**PS/VWF complex ELISA**

A PS/VWF complex ELISA was developed, capturing with a polyclonal goat anti-human VWF (Enzyme Research Laboratories) and detecting with a polyclonal goat anti-human PS (Enzyme Research Laboratories). 96-well microplates were coated overnight with anti-VWF and blocked for 2h with 2.5% casein (Sigma Aldrich, St. Louis, MO, USA). Purified protein samples in HBSA buffer or pooled control plasma were added to the wells and incubated for the indicated times. Samples were prepared with or without vortexing (2,500rpm, 30sec) or the addition of 1mM hirudin, 5mM Gly-Pro-Arg-Pro peptide, and 5mM $CaCl_2$. After removal of the samples, wells were washed five times with HBSAT (20mM HEPES, 150mM NaCl, 1% BSA, 0.1% Tween20), incubated with anti-PS for 90min and washed five additional times. Bound antibody was detected by incubation with o-phenylenediamine dihydrochloride (5-15min), and reactions were stopped with $H_2SO_4$.

**Spike protein IgG measurement**

Antibodies against SARS-CoV-2 spike protein were measured using an ELISA based on that of the Krammer group.[2,3] Briefly, purified spike protein was synthesized by the UK Protein Core Facility utilizing plasmid provided by Florian Krammer. Nunc-Immulon plates were coated with spike protein (2μg/mL), washed, blocked, and heat-inactivated (56˚C, 1h) serum (1:$10^2$ to 1:$10^5$ dilutions) was added to wells (2h, RT). Plates were washed, and HRP-conjugated goat anti-human IgG antibody was added (1h, RT). Plates were washed, developed for 10min using OPD substrate with acid stop, and read on a SpectraMax Microplate reader at 490nm. Endpoint titer was determined as the last dilution producing a signal +3SD background.

**Plasma tissue factor activity**

TF activity was measured using a spectroscopic factor Xa generation assay, adapted from the protocol of Hisada and Mackman,[4] as described in[5]. Briefly, 50µL PPP was diluted with 1mL hepes-buffered saline containing albumin (HBSA) (20mM HEPES, 150mM NaCl, 0.1% bovine serum albumin [BSA], pH 7.4) and centrifuged (20,000x*g*, 45min, 4°C) to isolate TF. The pellet was washed by centrifugation with HBSA and resuspended in 100µL HBSA. FXa generation was measured immediately following isolation. 50µL of each sample was incubated with FVIIa (12.5nM) and FX (375nM) in HBSA with 10mM $CaCl_2$, and Spectrozyme FXa (0.5mM). The cleavage of Spectrozyme FXa was monitored at 405nm for 2h at 37°C and compared to a TF standard curve to calculate TF activity, using nonlinear regression (GraphPad Prism v.8.0.2). Results were normalized to plasma total protein.

**Thioflavin T staining**

Thioflavin T fluorescent staining was utilized to measure fibrin amyloid microclots in PPP, using protocol adapted from Pretorius, et al,[6] and modified to run in a microplate format. Briefly, fibrinogen-immunodepleted plasma or PPP samples were incubated with 5µM Thioflavin T for 30min at RT, protected from light. Five microliters of samples were then diluted with HBS-PEG (20mM HEPES, 150mM NaCl, 0.1% PEG8000, pH 7.4) to 50µL and plated onto a 96-well plate. Total fluorescence at the excitation wavelength of 450±10nm and the emission at 482±10nm was then measured using the BioTek Cytation 5 fluorescence plate reader and the Gen5 software (Agilent Technologies, Santa Clara, CA, USA). Results were normalized to the fluorescence value of fibrinogen-immunodepleted plasma as negative control and to total protein. All samples were measured in triplicates.

**Plasma thrombin generation**

Thrombin generation was measured with PPP using a modified Calibrated Automated Thrombography to measure APC/PS pathway activation using thrombomodulin supplementation, as previously described,[5] in patient plasmas or control plasmas (pooled healthy normal, PS- and VWF-immunodepleted, and VWD type 3 deficient plasma). Briefly, 40µL of samples was incubated with 10µL of thrombin calibrator; and 10µL reagents containing 4µM final concentration of phospholipid vesicles (-TF), phospholipid vesicles and 1pM TF (-TM), or phospholipid vesicles, TF and 20nM thrombomodulin (+TM). PS activity was also assayed using APC supplementation, adapted from Brugge et al.,[7] in control plasmas. Here, 40µL of samples was incubated with 10µL of thrombin calibrator; and 10µL reagents containing 30µM PL, 6.8pM TF, with and without 5nM APC. Samples were incubated for 10min at 37°C, and thrombin generation was initiated with the addition of 10µL of a mixture of calcium and fluorogenic thrombin substrate. Thrombin activity was measured using a Fluoroskan Ascent Microplate Reader (Thermo Scientific) and quantified using Thrombinoscope software (Diagnostica Stago). Results (peak thrombin or endogenous thrombin potential (ETP)) were expressed as a ratio (with TM divided by without TM to quantify APC/PS activity, or with APC divided by without APC to quantify PS activity).

**Crosslinking and SA pulldown**

Plasma crosslinking and label transfer experiments were performed using Sulfo-SBED Biotin Label Transfer reagent (Thermo Scientific) per manufacturer's recommended protocol. Briefly, sulfo-SBED-labeled purified PS (25µg/mL) or VWF (10µg/mL) were added into 1mL of healthy normal plasma (Siemens Healthineers) and either PS-immunodepleted or VWD type 3 deficient plasma, respectively. Plasmas were then incubated (30min, RT, protected from light) with or without shear by vortexing, and then irradiated (365nm, 1hr, on ice) with UV lamp. Streptavidin agarose resin beads (Thermo Scientific) were washed 2x in PBS and 250µL were added into each (1mL) samples and incubated at RT with shaking. Beads were isolated with spin columns (1000xg for 1min) and washed 5x with PBS. Elution of biotinylated proteins were according to Cheah et al.[8] Briefly, beads were recovered into microcentrifuge in elution buffer (25mM biotin and 0.4% SDS in PBS) and boiled in 95°C for 5min. Samples were then centrifuged (1000xg for 2min) to get the supernatant, which was boiled with SDS sample buffer (95°C for 2min). SDS-PAGE was then performed as described.

**Native gel electrophoresis and immunoblotting**

Human protein S (HPS; Enzyme Research Laboratories, IN, USA) and plasma in native sample buffer were subjected to SDS-PAGE on 4-20% Mini-PROTEAN TGX precast protein gel (BioRad, Laboratories, CA, USA) and transferred to nitrocellulose membrane, using a Mini-PROTEAN Tetra Cell system (Bio-Rad) per manufacturer recommendations. Immunoblotting was performed using 2µg/mL sheep anti-human PS (CL20105A; Cedarlane Laboratories, Burlington, Canada), sheep anti-human PC (PAHPC-S; Haematologic Technologies, Essex Junction, VT, USA), mouse anti-human Mer (MAB8911; R&D Systems, MN, USA), rabbit anti-human-C4BP-β (Abcepta, San Diego, CA, USA), rabbit anti-human-TFPI (ProteinTech, Rosemont, IL, USA), mouse-anti-human-FV HC+LC monoclonal antibodies (Haematologic Technologies, Essex Junction, VT, USA), or 1µg/mL sheep anti-human VWF (Abcam, Cambridge, UK) primary antibodies, with 0.5µg/mL HRP-conjugated donkey anti-sheep IgG (Jackson ImmunoResearch, West Grove, PA, USA), horse anti-mouse IgG (Vector Laboratories,

Newark, CA, USA), and goat anti-rabbit IgG (Vector Laboratories, Newark, CA, USA) secondary antibodies.

**VWF multimer analysis**

Plasma VWF multimer distribution was assayed by SDS-agarose gel electrophoresis and immunoblotting, adapted from protocols by Thomazini, et al.[9] and Ott, et al.[10]. Electrophoresis and membrane transfer were using the Mini-PROTEAN Tetra electrophoresis system and Mini Trans-Blot electrophoretic transfer cell (Bio-Rad, Hercules, CA, USA). Briefly, buffers for SDS-agarose gel electrophoresis were prepared for a 0.8-1.6% (w/v; stacking-resolving) discontinuous gel electrophoresis using Seakem HGT Agarose (Lonza, Basel, Switzerland). Gel glass cassette and stand, gel comb, and serological pipet were prewarmed (50ºC, 15min) prior to pouring of the microwave-heated gel solutions into the gel cassette, then the comb was inserted onto the stacking gel, and maintained at RT until solidification. Agarose gels were then kept at 4ºC for 30min. Plasma samples were normalized by the VWF:Ag ELISA results with agarose sample buffer to reach a final 1µg/mL in 10µL loading per well, whereas purified PS was 25ng and VWF 10ng. Shearing of samples were performed by vortexing at ~2,500rpm for 30s. Recalcification of plasma entails the addition of 1mM hirudin (Sigma Aldrich, St. Louis, MO, USA), GPRP peptide (GenScript, Piscataway, New Jersey, USA), and 5mM $CaCl_2$.

Freshly prepared samples were heated at 56ºC for 30min prior to loading. Electrophoresis of the samples was then conducted at 10mA constant current at 9ºC using precooled electrode buffer until the dye front had migrated to the bottom of the gels. To increase transfer efficiency of the higher multimers, the gel was incubated in 1mM β-mercaptoethanol in transfer buffer for 30min. Transfer to nitrocellulose membrane was conducted at constant 120V and 4ºC for 1h. Immunoblotting was performed with either the LiCor system with IRdye antibodies (LiCor

Biosciences, Lincoln, NE, USA) or the BioRad enhanced chemiluminescence with horse radish peroxidase-conjugated antibodies. Primary antibodies used were sheep anti-human PS (Cedarlane Laboratories, Burlington, Canada), mouse anti-human-VWF-A2 (R&D Systems) at 2μg/mL, and rabbit anti-human C4BP-α (LifeSpan Biosciences, Seattle, WA, USA). Secondary antibodies were IRdye 800CW donkey anti-goat IgG, IRdye 680LT donkey anti-mouse IgG, and IRdye 800CW donkey anti-rabbit IgG (LiCor Biosciences, Lincoln, NE, USA) at 0.1μg/mL, and HRP-conjugated donkey anti-sheep IgG (Jackson ImmunoResearch, PA, USA) and horse anti-mouse IgG (Vector Laboratories, CA, USA) at 0.5μg/mL. Images were taken using either Image Studio (LiCor Biosciences) or ChemiDoc MP (BioRad).

**VWF cleavage by rADAMTS13**

VWF cleavage of recombinant ADAMTS13 was assayed with constant shearing (~2,500rpm) at RT for 60m with a tabletop mixer followed by immunoblotting, following protocol adapted from Han, et al.[11] Briefly, for each well, 1μL of control plasma N was diluted to 20μL and supplemented with 150nM purified VWF, 5mM GPRP peptide, 2mM AEBSF (Thermo Scientific, Waltham, MA, USA), 5mM $CaCl_2$, with or without 20mM EDTA, 50nM recombinant ADAMTS13 (Novus Biologicals, Centennial, CO, USA), or 150nM PS, in 0.2 mL PCR tubes. Some samples were then sheared for 60min using Retsch MM301 mixer mill (Glen Mills, Clifton, NJ, USA), mixed with agarose sample buffer, heated at 56ºC for 30min, and SDS-agarose electrophoresis, transfer, and immunoblotting performed according to previously described procedure.

**Bioflux microfluidics**

Microfluidic experiments were conducted using the BioFlux 200 (Fluxion Biosciences). Briefly, the Bioflux plate was coated with 40μg/mL collagen for 1hr at RT, before blocking with PBS supplemented with 0.5% serum albumin. Freshly drawn whole blood was supplemented with 10µg/mL AlexaFluor555-conjugated VWF and 200nM AlexaFluor488-conjugated PS, either with or without prior shearing at ~2,500rpm for 30sec. Samples were then anticoagulated (4U/mL hirudin), recalcified (1mM $CaCl_2$), and perfused at 35dyne/$cm^2$. After perfusing, channels were washed with PBS/0.5% BSA, and images were taken with a Nikon Eclipse T*i*2 microscope.

**Disrupted-flow microfluidics**

We visualized PS binding to self-associated VWF using a polydimethylsiloxane (PDMS) microfluidic device, with a block in the center of the channel, as previously described.[24] VWF was diluted in Tris-buffered saline (TBS, with or without 2mM $CaCl_2$) to 7.5µg/mL and was perfused at a flow rate of 0.02mL/min, resulting in VWF self-association over the PDMS block. PS (80% unlabeled, 20% labeled with Alexa Fluor-488) was diluted to 7.5µg/mL and perfused through the channel. VWF self-association was visualized with DIC and PS binding was visualized with epifluorescence microscopy.

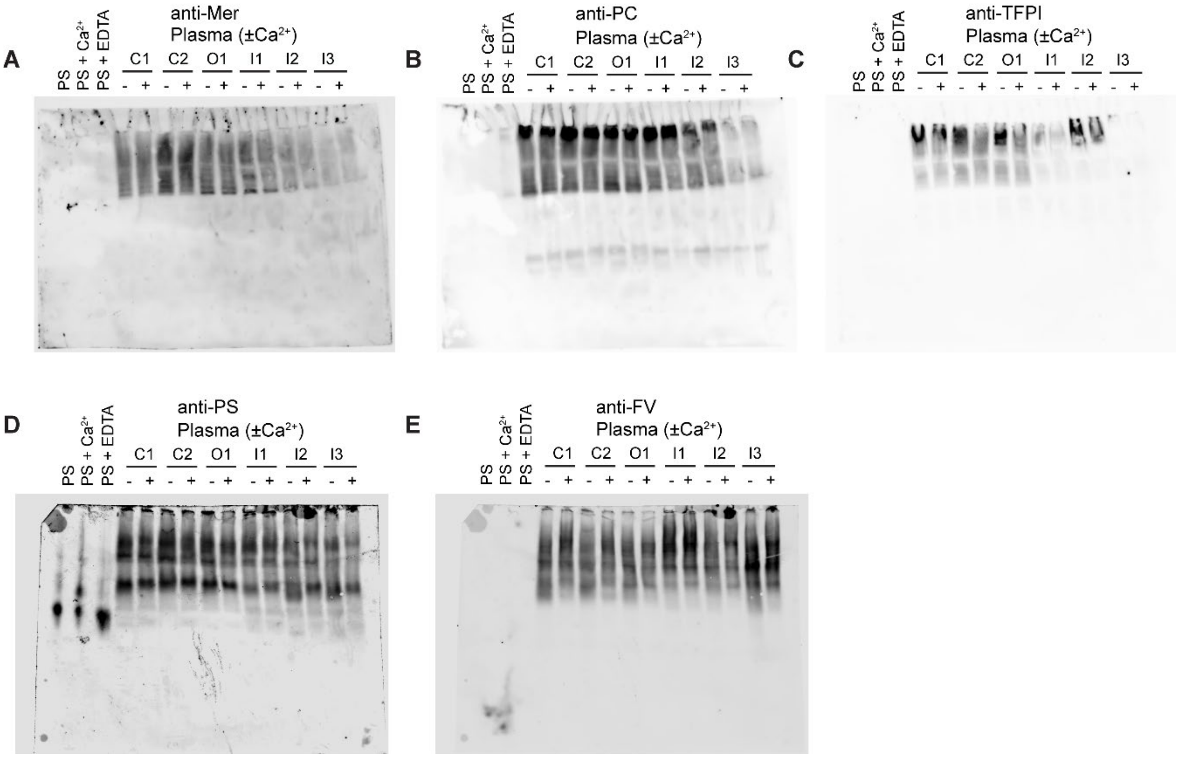


**Figure S1. Native gel electrophoresis shows that PS has many binding partners in plasma.** Purified PS (50ng), with either 20mM $CaCl_2$ or 5mM EDTA, or 4µL citrated plasma (C1-2: healthy controls, O1: COVID-19 outpatient, I1-3: COVID-19 inpatients), with or without 1µM Hirudin, 5mM GPRP peptide, and 5mM $CaCl_2$, were separated with native gel electrophoresis and probed with **(A)** anti-Mer, **(B)** anti-protein C, and **(C)** anti-TFPI antibody. A separate blot was probed with **(D)** anti-PS and **(E)** anti-factor V. (*, apparent monomer band; **, apparent dimer band)

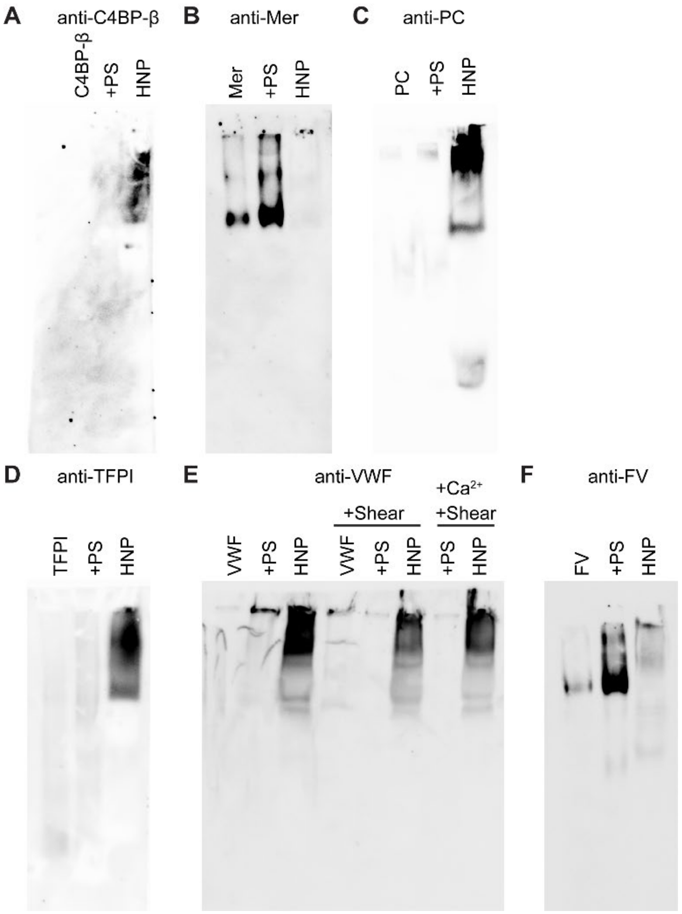


**Figure S2. Native gel electrophoresis of individual purified proteins.** Physiological concentration of each protein (200nM C4BP, 4nM Mer, 70nM PC, 5nM TFPI, 38nM VWF, 30nM FV) equivalent to 4µL of healthy normal plasma (HNP), in the absence or presence of 400nM PS, or 4µL HNP, were separated with native gel electrophoresis and probed with **(A)** anti-C4BP-β, **(B)** anti-Mer, **(C)** anti-PC, **(D)** anti-TFPI, **(E)** anti-VWF, **(F)** anti-FV antibody. Shearing entails vortexing at ~2,500rpm for 30sec and calcium entails supplementation with 1µM Hirudin, 5mM GPRP peptide, and 5mM $CaCl_2$.

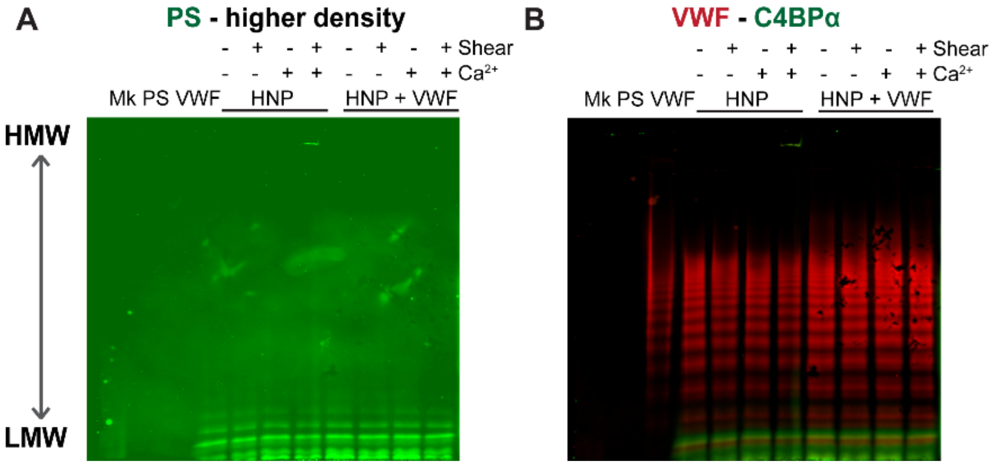


**Figures S3. PS co-migrates with lower molecular weight VWF multimers.** 25ng of PS or 10ng of VWF purified proteins, and 1µL of healthy normal plasma (HNP) from Siemens, with or without shearing (~2,500rpm vortexing for 30sec), or 1µM Hirudin, 5mM GPRP peptide, and 5mM $CaCl_2$ supplementation, or additional 10µg/mL of purified VWF, were separated for a VWF multimer analyses using SDS-agarose electrophoresis. The highest band on the molecular weight marker was 460kDa. **(A)** Shown is PS in green, tuned up in fluorescence intensity (800nm). **(B)** The same blot probed for VWF (red) and C4BP-α (green).

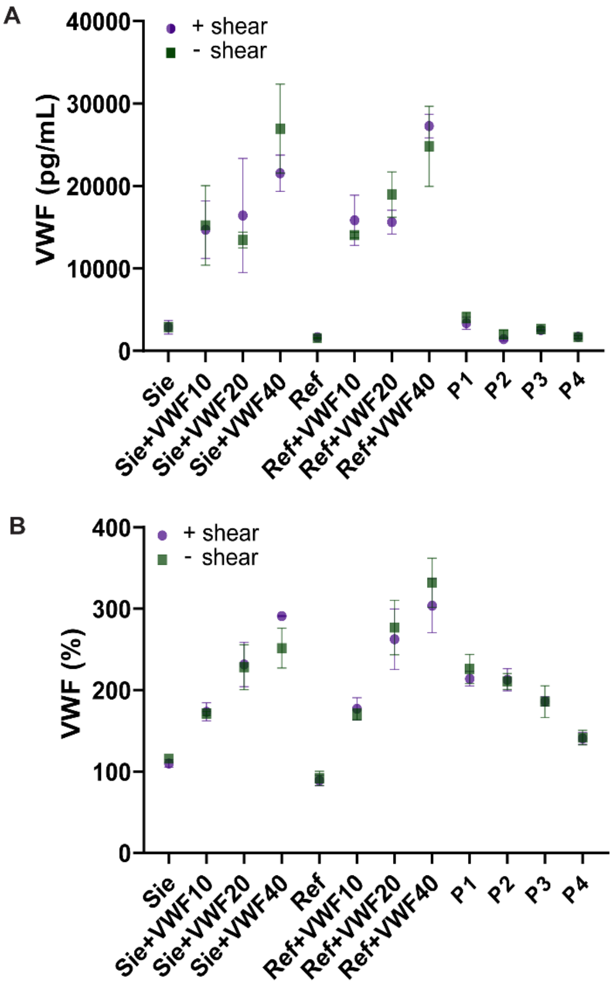


**Figure S4. Shearing does not interfere with VWF ELISA measurements.** VWF:Ag ELISAs (**A**, R&D systems; **B**, Enzyme Research Labs) results of healthy normal plasma (Sie, Siemens; Ref, Reference), with additions of purified VWF, and several COVID-19 in- (P1, P2) and outpatients (P3, P4), with or without shear.

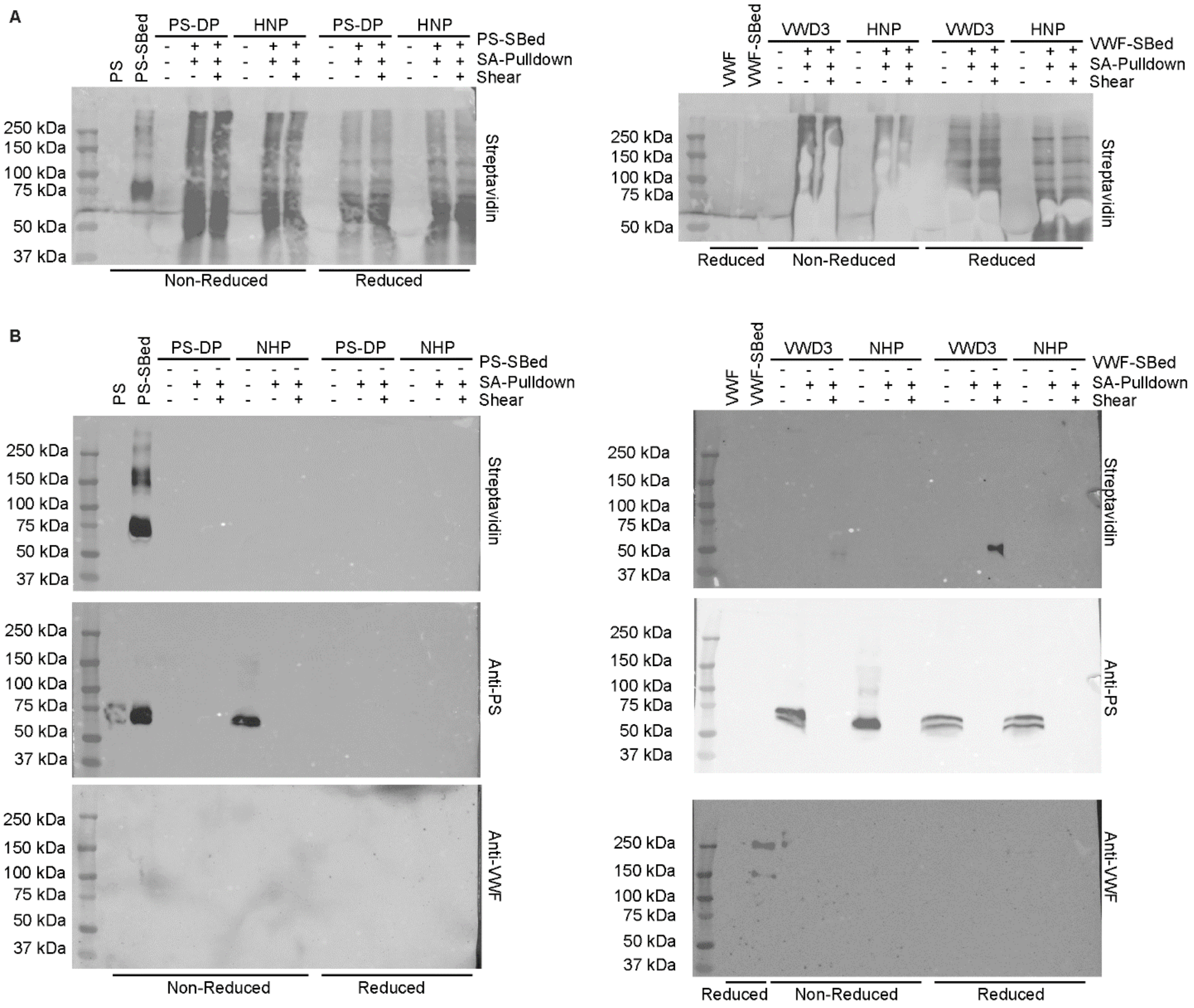


**Figure S5. PS and VWF do not non-specifically interact with streptavidin beads.** (**A**) Sulfo-SBED-labeled PS (25µg/mL) (left) or VWF (10µg/mL) (right) was incubated in PS-depleted plasma (PS-DP) or plasma from a patient with Type III VWD (VWD3), respectively, or in pooled healthy normal plasma (HNP), in the presence or absence of shearing. Crosslinking was induced by exposure to UV light (365nm), and biotinylated proteins were isolated using streptavidin-coated agarose beads. Eluted proteins were identified by blotting with Streptavidin-HRP. (**B**) Experiments were performed as in (A), using unlabeled PS (left) and VWF (right). Blots were probed used Streptavidin-HRP (top) or polyclonal antibodies against PS (middle) or VWF (bottom).

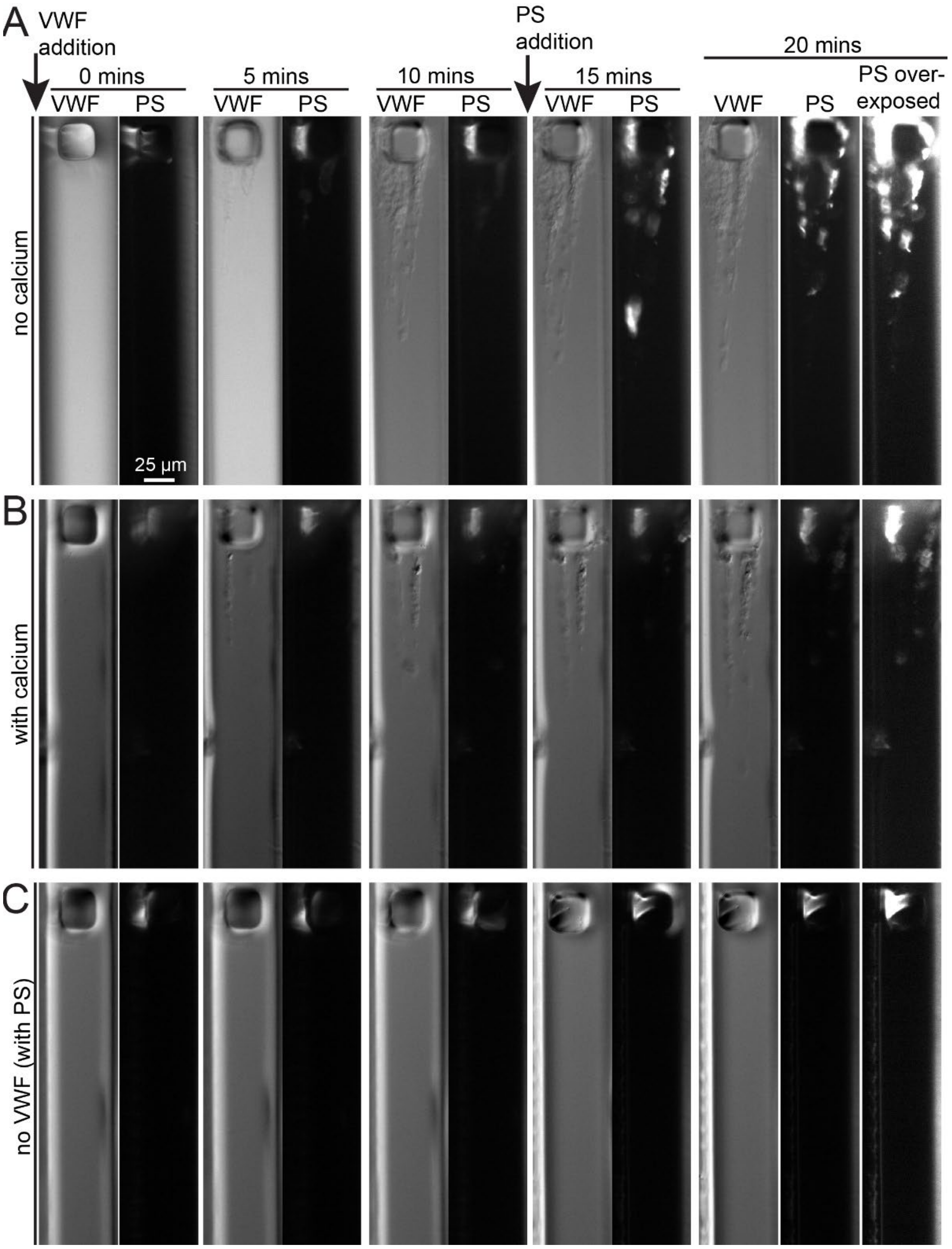


**Figure S6.** Representative images captured over time (0min, 5min, 10min, and 20min) of **(A)** VWF + PS without calcium, **(B)** VWF + PS with calcium (2mM $CaCl_2$), and **(C)** PS without VWF (and without calcium). VWF is added at time=0 and VWF self-association is visible around the block in the DIC images. At time=10min, fluorescently labeled PS is added and is observed binding to VWF, especially in the absence of calcium (A and B). Intensity in the fluorescent channel is not observed in the absence of VWF (C).

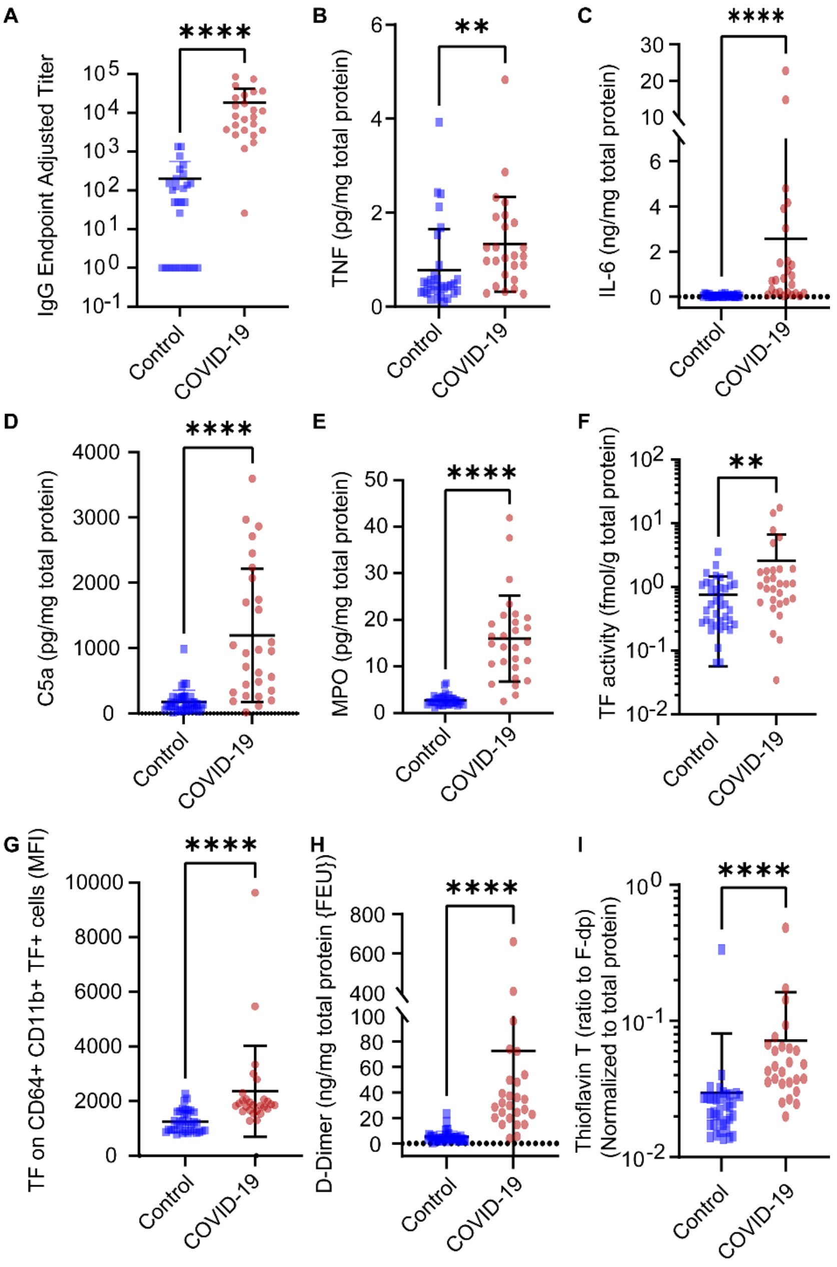


**Figure S7. COVID-19 inpatients show signs of inflammation associated thrombosis.** Citrated plasma samples and peripheral blood derived mononuclear cells were collected from healthy controls (n=38) and COVID-19 inpatients (n=30). Due to sample limitations, some measurements could not be performed with all subjects. **(A-E, H)** Shown are ELISA measurements of plasma **(A)** anti-spike protein IgG, **(B)** tumor necrosis factor (TNF), **(C)** interleukin-6 (IL-6), **(D)** Complement Component 5a (C5a), **(E)** Myeloperoxidase (MPO), and **(H)** D-dimer Fibrinogen Equivalent Units, **(F)** plasma tissue factor (TF) activity as measured by factor Xa activity assay, **(G)** monocyte TF expression as measured by flow cytometry, and **(I)** plasma thioflavin T fluorescence staining which measures fibrin as adapted from Pretorius et al.[6]

Blue squares represent controls and red circles represent COVID-19 inpatients. All data were normalized to total protein. Every data point is the average of three replicates (mean±SD). *P*-values are according to Mann-Whitney test (**, $P < 0.01$; ****, $P < 0.0001$).

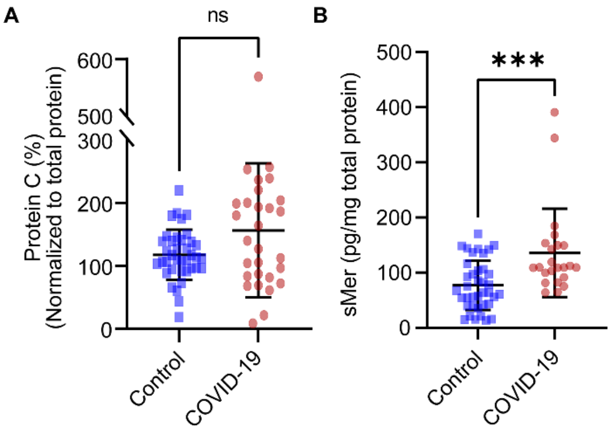


**Figure S8. Free protein S (PS) deficiency in COVID-19 patients is not explained by increases in known PS-binding proteins.** Citrated plasma samples were collected from healthy controls (n=38) and COVID-19 inpatients (n=30). Due to sample limitations, some measurements could not be performed with all subjects. Shown are ELISA measurements of **(A)** protein C and **(B)** shed Mer tyrosine kinase. All data were normalized to total protein. Every data point is the average of three replicates (mean±SD). Blue squares represent controls and red circles represent COVID-19 inpatients. *P*-values are according to Mann-Whitney test (n.s., non-significant; ***, $P$<0.001).

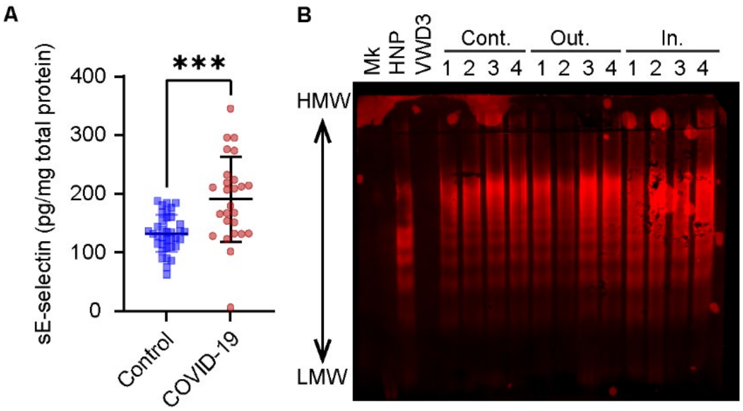


**Figure S9. Plasma von Willebrand Factor (VWF) is higher in patients with severe COVID-19. (A)** E-selectin was measured using ELISA in citrated plasma samples collected from controls (n=38) and COVID-19 inpatients (n=30). Every data point is the average of three replicates (mean±SD). *P*-values are according to Mann-Whitney test (***, *P*<0.001; ****, *P*<0.0001). **(B)** Healthy normal plasma (HNP) from Siemens and citrated plasmas from VWD type 3 patient (VWD3), healthy controls (Cont.; 1-4), COVID-19 outpatients (Out.; 1-4), and COVID-19 inpatients (In.; 1-4) were normalized to 10µg/mL of VWF:Ag, and 1µL of each samples were separated for VWF multimer analysis using SDS-agarose gel electrophoresis and anti-VWF probing.

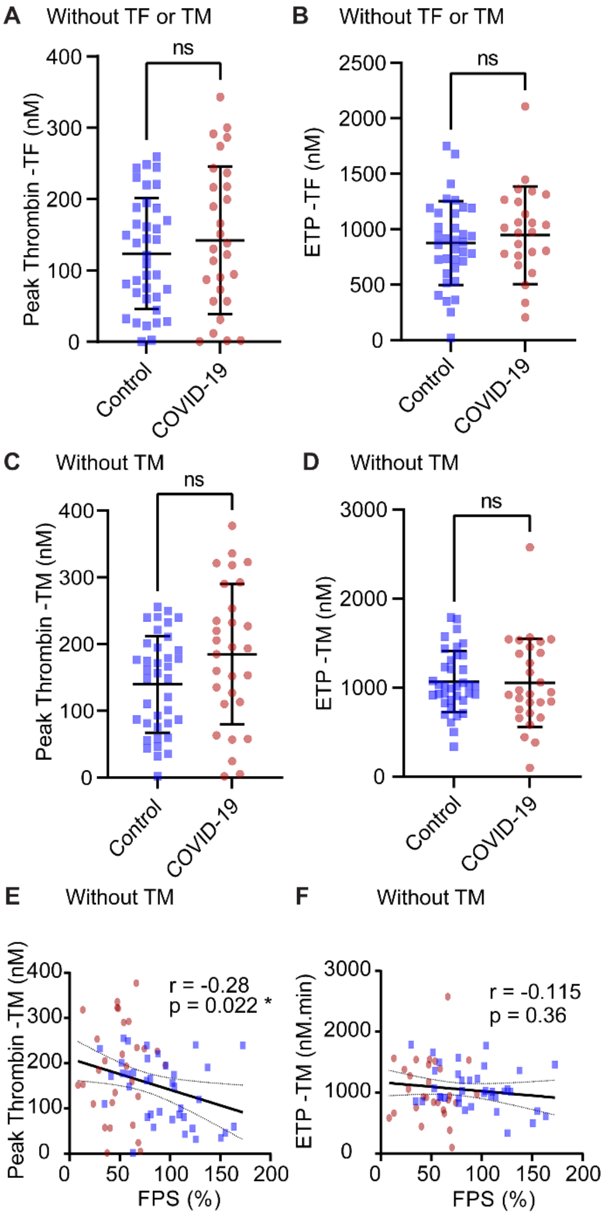


**Figure S10. Plasma thrombin generation in COVID-19 inpatients shows less APC/PS activity.** Plasma thrombin generation is measured using fluorogenic substrate to thrombin according to the Calibrated Automated Thrombography (Thrombinoscope) method (Diagnostica Stago) in citrated plasma samples collected from healthy controls (n=38) and COVID-19 inpatients (n=30). Several inpatients were receiving heparin prophylactic dose at the time of blood collection. Shown are **(A, C)** peak thrombin concentrations and **(B, D)** Endogenous Thrombin Potential (ETP) in assays initiated with 4µM phospholipids **(A-B)** without and (**C-D**) with 1pM tissue factor (TF), *P*-values are according to Mann-Whitney test (n.s., non-significant); **(E-F)** Free PS plotted against **(E)** peak thrombin concentration and **(F)** ETP in assays without thrombomodulin supplementation, *P*-values and r correlation coefficients are according to Spearman correlation analysis (n.s., non-significant; *, $P<0.05$). Every data point is the average of three replicates (mean±SD). Blue squares represent controls and red circles represent COVID-19 inpatients.
